# Acquisition of an immunosuppressive microenvironment after CAR-T therapy drives T-cell dysfunction and resistance

**DOI:** 10.1101/2024.12.20.629119

**Authors:** Marianna Ponzo, Lorenzo Drufuca, Chiara Buracchi, Marco M. Sindoni, Silvia Nucera, Cristina Bugarin, Ramona Bason, Grazisa Rossetti, Raoul Bonnal, Cristian Meli, Benedetta Rambaldi, Federico Lussana, Silvia Ferrari, Alex Moretti, Giulia Risca, Christian Pellegrino, Markus G. Manz, Stefania Galimberti, Alessandro Rambaldi, Giuseppe Gaipa, Andrea Biondi, Massimiliano Pagani, Chiara F. Magnani

## Abstract

Chimeric antigen receptor (CAR) T cells targeting CD19 induce durable responses in B-cell acute lymphoblastic leukemia (B-ALL). However, the contribution of the tumor microenvironment to the therapeutic response after CAR-T cell treatment remains incompletely understood. We performed single-cell RNA sequencing and spectral flow cytometry-based analyses of bone marrow-resident immune cells from B-ALL patients before and after CAR T-cell treatment. We observed profound changes in the microenvironment in response to CAR T cell-mediated inflammation, including an increase in myeloid cells. Significant induction of the IFN response, hypoxia, and TGF-β-signaling was associated with expansion of myeloid-derived suppressor cells (MDSCs) and endogenous exhausted CD8+ T cells. PD1 expression in endogenous T cells post-treatment was associated with a lack of durable response in the cohort of patients analyzed. Further, we revealed that HIF1α, VEGF, and TGFBR2 are key players in the intercellular communication between CAR T cells and the immune niche, driving widespread T-cell dysfunction. Infusion of anti-CD19 CAR T cells led to increased accumulation of human MDSCs, exacerbation of a hypoxic environment and T-cell exhaustion in HSPC-humanized mice bearing a human tumor. In conclusion, CAR T-cell-mediated myeloid activation is associated with pathways of immune dysregulation that may antagonize the effects of therapy.

**Key points:**

- The tumor immune context post-treatment affects CAR T-cell fate and endogenous immunity in B-cell acute lymphoblastic leukemia.
- IFN response, hypoxia, and TGF-β-signaling result in endogenous T-cell exhaustion and compromise CAR T-cell efficacy.

## Introduction

Chimeric Antigen Receptor (CAR) T cell-based therapies targeting the CD19 antigen have been approved for the treatment of children and adults with relapsed/refractory (R/R) B cell acute lymphoblastic leukemia (B-ALL), based on the results achieved in the pivotal Eliana and ZUMA-3 studies.^1,2^ To reduce costs and regulatory requirements associated with CAR T-cell manufacturing,^3^ our group generated anti-CD19 CAR T cells with the non-viral Sleeping Beauty transposon vector from donor T cells (CARCIK-CD19), which proved potent anti-leukemic activity associated with the absence of severe toxicities.^4^ Despite these promising results, only 40-50% of B-ALL patients treated with any type of CAR T cells maintained a durable remission long term.^2,5^ In a considerable portion of treated patients, relapse occurs in the presence of CD19-expressing tumors, mainly because of CAR T-cell dysfunction.^1,2,5^

B-ALL is a disease that mainly resides in the bone marrow (BM) and CAR T cells do not act on cancer cells in isolation, but within the tumor microenvironment (TME), which is known to affect T-cell activity.^6^ Therefore, a major interest is to identify factors within the TME that influence the activity and potency of anti-CD19 CAR T cells in B-ALL. B-ALL TME protects leukemia^7^ and, conversely, suppresses effector T cells.^8^ However, the interaction between the BM microenvironment and CAR T cells in B-ALL has not been investigated. Moreover, the respective role of the immune landscape in affecting the therapeutic efficacy remains undefined.

We hypothesized that the immunological niche of the BM reacts to CAR T cell-mediated inflammation by modulating the immune response through the activation of inhibitory pathways and molecules. To test our hypothesis, we performed single-cell transcriptional analysis (scRNA-seq) and subsequent validation using a spectral flow cytometer of BM samples from anti-CD19 CAR T cells and CARCIK-CD19 treated patients. We identified hypoxia, VEGF, IFN, and TGF-β pathways in association with the expansion of myeloid-derived suppressor cells (MDSCs) and exhaustion of resident CD8+ T cells and infused CAR T cells. Further investigation with leukemia-bearing hematopoietic stem/progenitor cell (HSPC)-humanized mouse models recapitulated the human TME and the observed immunosuppressive response post-treatment. These data suggest a role of the B-ALL TME in inducing T-cell dysfunction in response to CAR T-cell treatment, with the potential to antagonize the effect of the therapy.

## Methods

Detailed information is provided in the Supplementary Materials file (available on the Blood Web site).

### Patient Samples

Data in this study were generated from patients enrolled in the FT01CARCIK Phase I/IIb clinical trial (NCT03389035), in the FT03CARCIK Phase II study (NCT05252403) or from autologous CAR T cells in the context of either phase I/II trial or commercial cell therapy (tisagenlecleucel).

### scRNA-seq

The scRNA-seq libraries were generated using Chromium Single Cell 5′Reagent Kit of the 10x Genomics (v3.1). Non-PCR duplicates with valid barcodes and Unique molecular identifiers (UMIs) were employed to create the gene-barcode matrix according to standard pipelines from the Cell Ranger software (Chromium Single Cell Software Suite).

### Flow Cytometry Validation

BM samples from treated patients were thawed and stained with a 30-colors panel. Data were acquired on a Cytek Aurora (Cytek® Biosciences) and analyses were performed using Infinicyt (Cytognos), DIVA software, and FlowSOM.

### In vivo studies

NSG (NOD.Cg-Prkdcscid Il2rgtm1Wjl/SzJ) mice were humanized as previously reported.^9^ n-hu-Nalm-NSG mice were generated by transplanting Nalm6 after the establishment of the human hematopoiesis.^10^ n-hu-PDX-NSG mice were generated by sub-lethal irradiation of newborn mice and co-transplantation of healthy CD34+ HSPCs from mobilized peripheral blood cells and patient-derived xenograft cells (Patient ID #003).^11^

## Results

### CAR T cells elicit an acute response involving both innate and adaptive immunity

To assess how CAR T cells influence the kinetics of the immune reconstitution following lymphodepletion, we analyzed CAR T cells and total CD3+ T cells by flow cytometry in parallel to common hematology laboratory parameters in the peripheral blood (PB) of 15 patients with B-ALL. Patient characteristics are reported in STable 1. CAR T-cell engraftment was detected with a median time to maximal expansion of 14 days (range 10-28 days) and a median peak expansion of 56 cells/mmc (range, 2-887), followed by a decrease in the third and fourth weeks post-infusion. A similar dynamic of expansion and contraction was observed in endogenous CD3+ T cells, with a median peak expansion 836 cells/mmc (range, 68-4395) at day 21 (range, 7-28, Figure 1A). Interestingly, CD3+ T-cell kinetics appeared biphasic, as they experienced a contraction around 10 days post-infusion due to pre-treatment lymphodepletion, followed by a rapid recovery associated with, but lagging behind, CAR T-cell expansion, before declining thereafter. Multiphasic kinetics characterized by pre-treatment lymphodepletion-induced reduction, expansion and contraction dynamics were also observed for the absolute white blood cell count (WBC), lymphocyte count (ALC), monocyte count (AMC), and neutrophil count (ANC) with a median maximal expansion of 3890 cells/mmc (range, 240-8810), 1270 cell/mmc (range, 150-7390), 725 cell/mmc (range, 80-1200), 2763 cell/mmc (range, 30-4580), respectively (Figure 1B-C). This effect can only partially be explained by reconstitution after lymphodepletion, as it appears to be very different in terms of intensity and kinetics in similarly lymphodepleted patients, in whom the lymphocyte count does not change substantially from day 14 onward.^12^ This innate and adaptive response mirrored, albeit delayed, CAR T-cell expansion and contraction, suggesting that CAR T-cell-driven inflammation is actively regulated by a process of immune regulation (Figure 1D).

**Figure 1.**
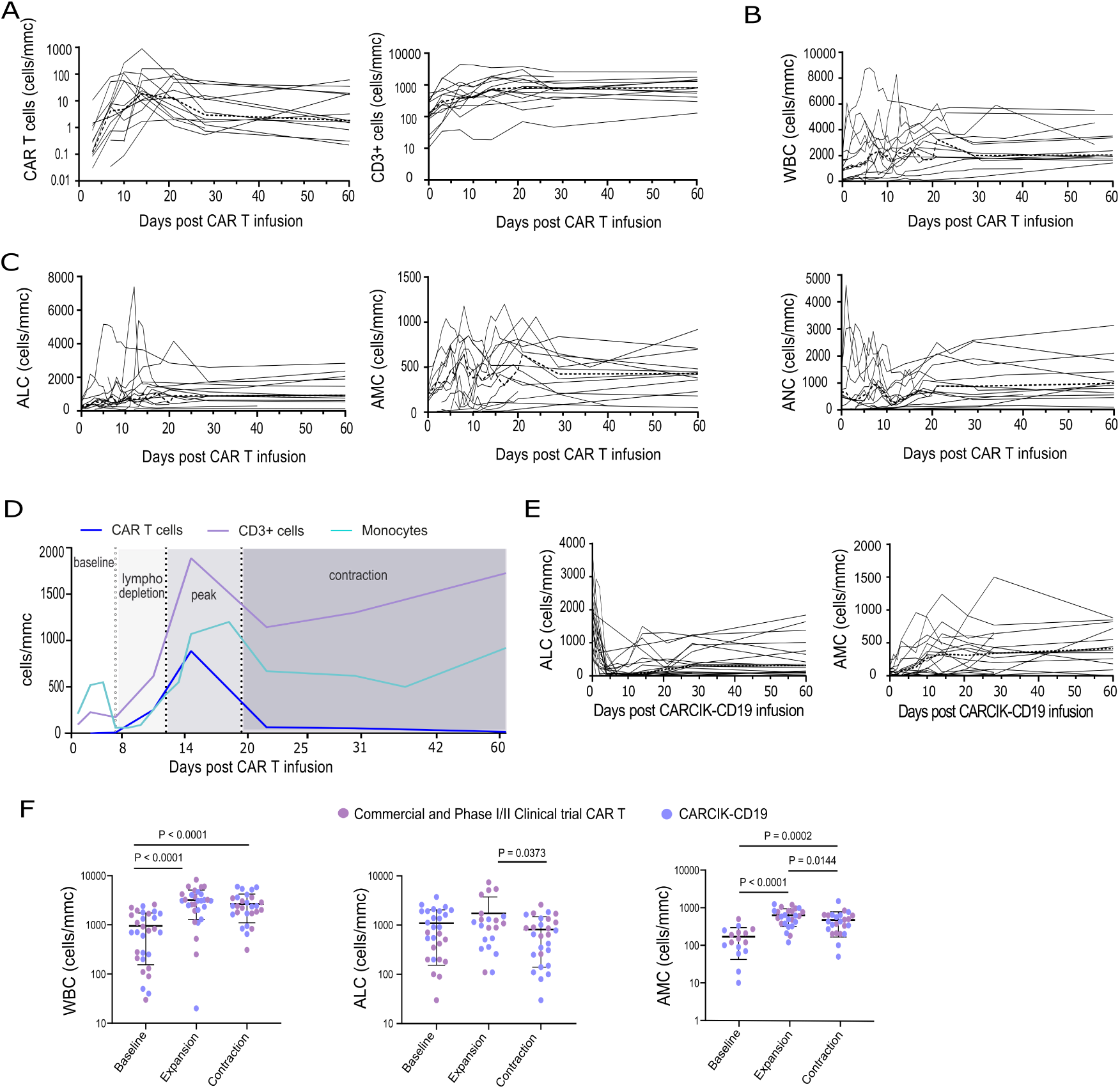
Multiphasic cellular kinetics after CAR T-cell infusion. (A) Absolute cell counts of CAR T cells and total CD3+ cells by flow cytometry in pediatric patients treated with autologous commercial and phase I/II anti-CD19 CAR T cells (n = 15). The median is depicted as a dashed line. (B) Absolute cell counts of total white blood cells (WBC). (C) Absolute lymphocyte count (ALC), absolute monocyte count (AMC), and absolute neutrophil count (ANC). (D) CAR T-cell, CD3+ cell and monocyte cellular dynamic of a representative pediatric B-ALL patient, showing a biphasic kinetic due to lymphodepletion, expansion and contraction. (E) ALC and AMC in patients treated with CARCIK-CD19 (n= 21). (F) WBC, ALC and AMC at baseline (pre-infusion), peak of expansion, and contraction (day 28 post-infusion). Colors indicate the type of IP. Data illustrate the mean ± SD, Statistical analysis performed using one-way ANOVA with Tukey’s multiple comparison.

A similar acute response of ALC (525 cell/mmc, range, 110-1500), AMC (490 cell/mmc, range, 20-1500), WBC (3320 cell/mmc, range, 380-7820), and ANC (1560 cell/mmc, range 80-5130), albeit less pronounced, followed by contraction, was confirmed in 21 patients treated with CARCIK-CD19 (Figure 1E and SFigure 1A-B). At day 28, a significant decrease in ALC and AMC was observed compared to the peak of expansion (Figure 1F, n=32 patients). Overall, these data suggest that CAR T cells elicit a bystander activation of endogenous innate and adaptive immunity. A widespread resolution of CAR T-driven inflammation occurs during the first month after infusion. Interestingly, the intensity of the inflammatory response seems to vary depending on the cell product used, as evidenced by inferior WBC, ALC and AMC expansion in CARCIK-CD19 compared to commercial and phase I/II autologous CAR T-cell infusion.

### scRNA-seq of the BM microenvironment after CAR T-cell infusion reveals an increase in myeloid cells

Having observed a contraction of the responses elicited by CAR T-cell infusion, we hypothesized that the TME was reacting to CAR T-cell mediated inflammation by accumulation of immune suppressor cells and induction of inhibitory pathways and molecules. To elucidate the involved cellular and molecular mediators, we performed scRNA-seq of BM samples from B-ALL patients collected at early time points (1-2 months) after treatment. CD45+CD3+ T cells, and CD45+/lowCD3-, encompassing the hematopoietic immune niche of the TME, were sorted and compared to cells of the corresponding patient’s BM before CAR T-cell treatment at the moment of relapse. In parallel, manufactured CAR T-cell infusion products (IP) were sorted according to surface CAR expression (Figure 2A-B).

**Figure 2.**
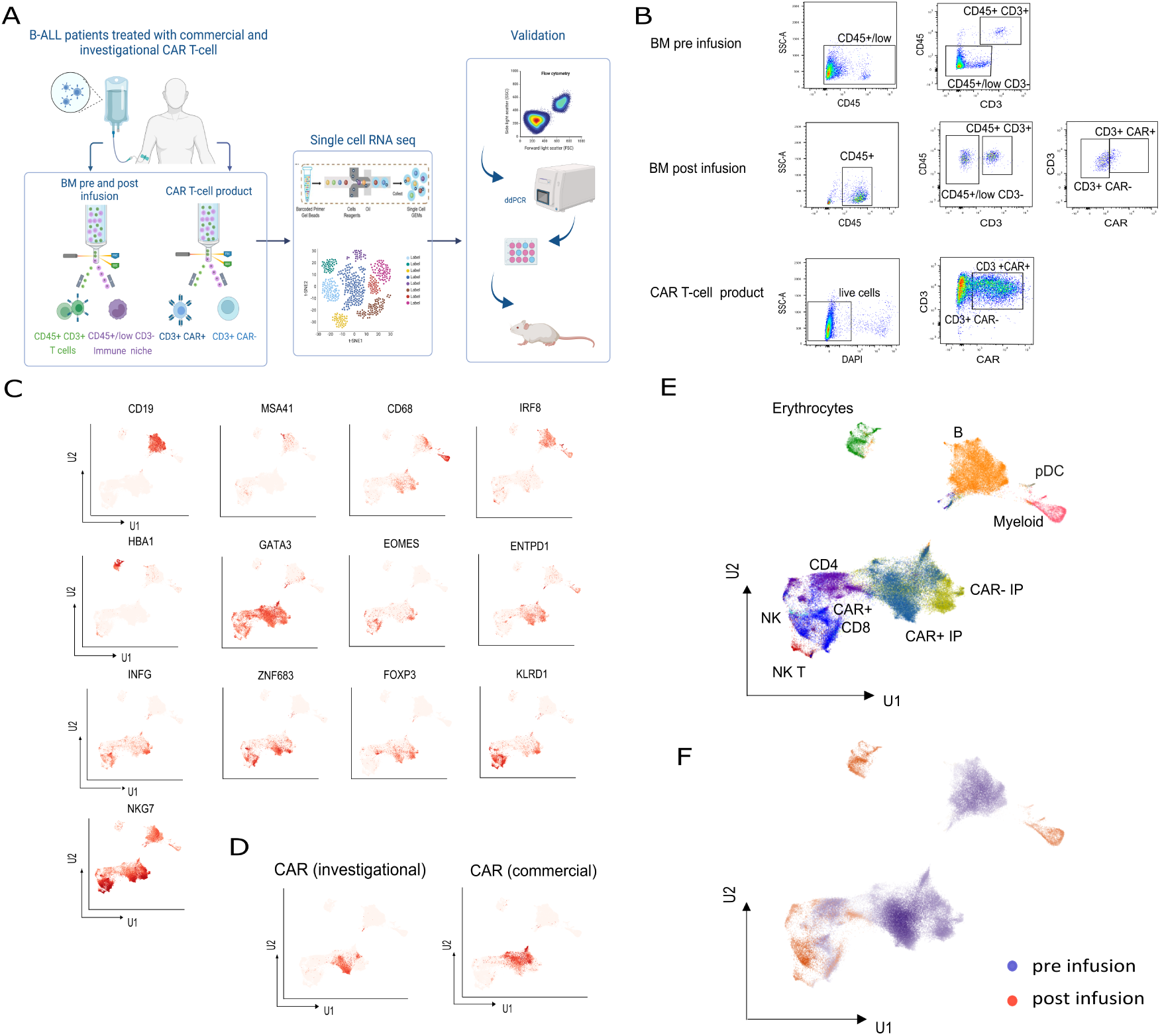
Single-cell RNA sequencing unveils profound restyling in the TME. (A) Experimental design and sorting strategy. Sequencing libraries have been generated from BM samples of pediatric B-ALL patients before and after anti-CD19 CAR T cell infusion, and the IP. Cells were sorted and subjected to scRNA-seq. Data were subsequently validated using spectral flow cytometry, ddPCR, in vitro co-cuture and in vivo humanized models. (B) Representative flow cytometry gating applied for sorting B-ALL BM samples and the IP. (C) Expression levels of lineage-specific genes overlaid on the Uniform Manifold Approximation and Projection (UMAP), (D) Transcript-based CAR T-cell cluster identification analysis. (E-F) UMAP embedding of multidimensional scRNA-seq data by cell clusters and time points.

Eighteen sequencing libraries were analyzed from three B-ALL patients (STable 1), for a total of 71.407 high quality cells at an average of 65.000 reads per cell (SFigure 2A and STable 2). One of these patients (#001) had been treated with CARCIK-CD19 (“investigational”) and two of them (#002 and #003) with commercial CAR T cells. All three patients achieved complete remissio<u>n.</u> However, patient #001 underwent consolidation with an allogeneic hematopoietic stem cell transplantation after achieving the remission for loss of B-cell aplasia, while the other two patients relapsed within 4 months after infusion. Extensive integration of the data coming from the individual patients was observed (SFigure 2B). Unbiased clustering was performed and 8 cell clusters were identified as CD4, CD8, NK, NKT, myeloid, B, plasmacytoid DC (pDC) and erythrocytes based on relative expression levels of lineage-specific genes (Figure 2C).^13^ To ensure that CAR+ T cells could be traced in post-infusion samples, we evaluated the sensitivity of CAR+ T-cell detection in complex cellular mixtures using an in silico limiting dilution assay (SFigure 2C). CAR gene expression was detected in all post-infusion samples. We applied high-resolution clustering of CD3+ cells, to recognize a qualitatively enriched cluster in CAR+ cells and annotate it as putative CAR+ (Figure 2D and SFigure 2D). The identified 11 subpopulations were visualized using UMAP (Figure 2E).

We observed profound changes in the composition of the BM compartment in samples post CAR T-cell infusion compared to pre-infusion samples (Figure 2F). Remarkably, a complete disappearance of the B-cell associated cluster was observed after treatment, which correlates with the clinical data of the selected patients who achieved complete remission. Concomitantly, we noticed a relative increase in the fraction of myeloid cells post-treatment.

### The myeloid compartment is skewed towards MDSCs following CAR T-cell therapy

We evaluated the abundance of the 6 transcriptionally distinct cell clusters identified in the BM-derived CD45+/lowCD3- cell compartments comparing samples post-infusion to pre-infusion (Figure 3A). Identified CD19+ malignant B cells and CD20+ healthy B cells (Figure 3B and SFigure 3) were the most represented compartments pre-treatment,^14^ while erythrocytes, myeloid cells, encompassing DC and monocytes, NK, and pDC significantly increased post-treatment (Figure 3A and SFigure 4A-B). In the myeloid cluster, Gene Set Enrichment Analysis (GSEA) across individual cell types using the Hallmark gene set collection^15^ showed a significant enrichment of gene expression associated with IFN-α and IFN-γ response, IL-6/JAK/STAT3 signaling, inflammatory response, and hypoxia, associated with upregulation of *HIF1α* and its target gene, the angiogenic factor *VEGFA* (Figure 3C). Using metabolically scoped gene set collections manually curated from KEGG and Reactome databases,^16^ we observed increased lipid metabolism, steroid biosynthesis, and glycosaminoglycan biosynthesis (SFigure 4C-D), which are associated with immunosuppressive myeloid cells.^17^ Differential gene expression analysis revealed up-regulation of molecules that orchestrate myeloid-derived suppressor cell (MDSC) immunosuppressive functions, such as *TGFB1*, *ARG2*, *NOS1AP*, *ENTPD1* (CD39), *S100A9* and *S100A12* (Figure 3D). Moreover, *CCL2* and *TLR2*, which play a key role in MDSC migration,^18^ and DC dysfunction,^19^ respectively, were also up-regulated. We applied a specific signature for murine MDSCs by mapping the overexpressed genes to human orthologues using BioMart tools,^20^ and found that myeloid cells from samples post-treatment display a higher resemblance to MDSCs (Figure 3E).

**Figure 3.**
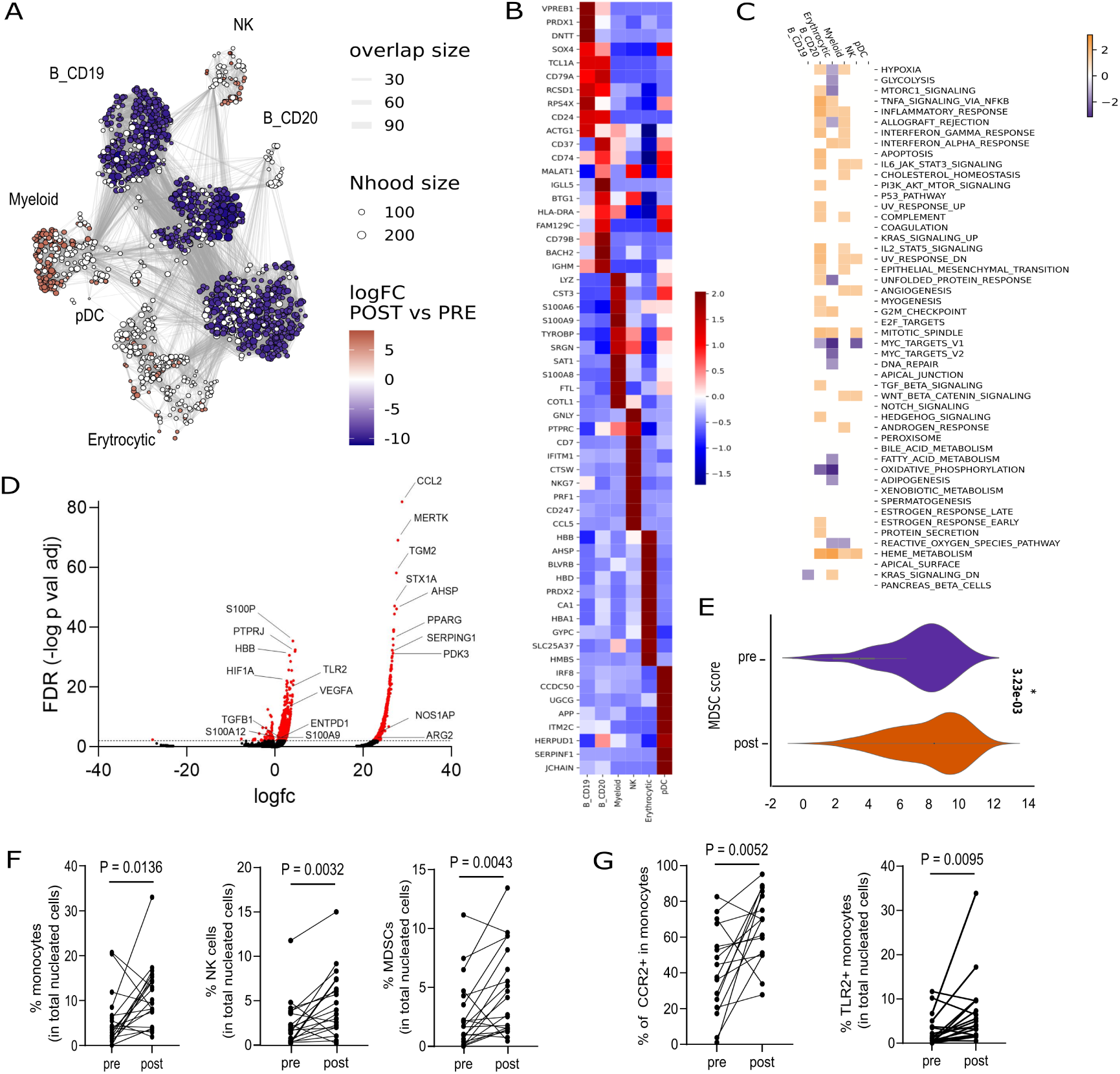
Bone Marrow Myeloid cells are enriched in Myeloid-Derived Suppressor Cells after CAR T-cell injection. (A) A neighborhood graph of the differential abundance test comparing samples post-infusion to pre-infusion with MILO and neighborhoods displaying significant abundance are expressed as log fold change (logFC). Size corresponds to the number of cells in each neighborhood (Nhood) in the UMAP space. Edge width (overlap size) is proportional to the number of cells shared between neighborhoods. As expected, limited overlap is observed between different cell types. (B) Gene expression Heatmap of the top ten cell-type-specific marker genes. (C) Significantly-enriched Molecular Signatures Database Hallmark gene sets in post-treatment versus pre-treatment within each immune cell types of the CD45+CD3- libraries (adjusted P val< 0.05). (D) Volcano Plots of DEG in myeloid cell cluster indicating genes up-regulated (rightward direction) and genes down-regulated (leftward direction) in post-CAR T-cell infusion compared to pre infusion. (E) MDSC signature scoring at different time points (arbitrary unit). (F) Flow cytometry analysis (n=20 paired pre- and post- treatment samples) with Infinicyt depicting frequencies of monocytes (CD33+CD14+ or CD16+), NK (CD3-CD56+) and MDSC (CD11b+CD33+HLA-DRlowCD14-CD15+ or CD14+CD15-) in total nucleated cells. Statistical analysis was performed using Wilcoxon-matched paired t-test. (G) Flow cytometry analysis depicting the frequencies of CCR2+ cells gated in monocytes and the frequencies of TLR2+ monocytes in total nucleated cells.

To validate and refine our findings, we expanded our study with a cohort of additional 20 matched BM samples (STable 1) pre- and post-CAR T-cell treatment acquired using spectral flow cytometry. Identification of NK and monocytes was based on reference guidelines by the EuroFlow Consortium using the Infinicyt software. MDSCs were identified according to their marker expression (CD11b+CD33+HLA-DR-/low CD14-CD15+ or CD14+CD15-, SFigure 5A). In line with our findings, we observed a significant increase in monocytes (median 3.7%, SD 5.8%, pre vs post 9.8%, SD 9.8%), NK (1.8%, SD 2.6% vs 3.7%, SD 3.6 %), and MDSCs (1.7%, SD 2.9%, vs 3.4%, SD 3.4%), complemented by a decrease in CD45low leukemic cells post-treatment (Figure 3F). To characterize specific treatment-related alterations in the myeloid compartment, we used manual gating. We found a significant increase of monocytes expressing high levels of CCR2, the receptor for CCL2, and TLR2 in total nucleated cells post-treatment, consistent with the scRNAseq data (Figure 3G and SFigure 5B). In sum, these data provide strong evidence for activation of signaling pathways in the TME that drive a skewing towards acquisition of an MDSC phenotype and immunosuppressive functions after CAR T-cell treatment.

### Increased T-cell exhaustion post-treatment affects both endogenous T cells and CAR T cells

Having shown a remodeling of the myeloid compartment after CAR T-cell infusion, we explored changes occurring in the CD3+ component to identify subsets of T lymphocytes associated with the therapy which may affect the therapeutic efficacy. We applied dimensionality reduction to the CD45+CD3+ scRNA-seq data using UMAP (Figure 4A) and utilized Garnett classification by integrating differential expression of cell-type specific genes and implementing a training of the classifier (SFigure 6A-B) to generate a final annotation (SFigure 6C-D).^21^ We identified 10 distinct clusters, including CD4 naïve-like, Th1/Th17, Th2, IL-10 producing Type 1 regulatory T cells (Tr1), Treg, CD8 effector, early memory, exhausted, resident memory, and NKT cells (Figure 4B). Global cell type enrichment analysis using SingleR transcription-based cell assignments revealed an increase in NKT, Tr1, and exhausted CD8+ T cells post CAR T-cell infusion (Figure 4C). A higher expression of genes associated with cell cycle in endogenous T cells, as well as with IFN-α and IFN-γ response and TGF-β signaling in CAR T cells was observed. Interestingly, we found widespread inhibition of glycolysis and oxidative phosphorylation (OXPHOS), which have been previously associated with metabolic insufficiency exhibited by exhausted T cells (Figure 4D).^22^ A shared signature of genes associated with lipid metabolism suggests that T cells sense a lipid-enriched and nutrient-deprived TME which have been reported to facilitate T-cell exhaustion (SFigure 7A-B).^23^ Endogenous exhausted T cells showed maintenance of some resident memory features,^24,25^ with a higher expression of *TGFB1, IL7R*, and *IL-32*, while CAR T cells up-regulated expression of *HIF1A*, *TOX*, *EOMES*, and *TGFB1* (Figure 4E). Interestingly, *TCF7,* which is critical in driving differentiation of progenitor exhausted cells,^26^ was up-regulated in CAR T cells as well as endogenous exhausted T cells. To validate this finding at a cellular and protein level, we analyzed the lymphoid cells from the flow cytometry data using the Infinicyt software and showed that the number of CD3+ and CD3+CD8+ cells significantly increased while CD3+CD4+ cells decreased after treatment. This was also evident when we performed hierarchical consensus clustering using FlowSOM (Figure 4F-G). In addition, we observed an increase of effector T cells and terminal effector CD3+CD8+ cells (SFigure 8A-B). We analyzed the expression of the exhaustion markers by manual gating and observed an increase in the % of CD3+PD1+ (median 7.50%, SD 7.7%, pre vs post 16.5%, SD 11.9%), CD3+TIGIT+ (4.9%, SD 7.3% vs 12.6%, SD 10%) and CD3+TIM3+ (0.7%, SD 2.4% vs 6.2%, SD 11.2%) as well as double-positive TIGIT+PD1+CD3+ T cells (Figure 4H and SFigure 8C-D). Notably, CAR T cells showed increased expression of TIGIT, PD1 and TIM3. PD1 expression was also increased in resident CD4+ cells post-infusion (Figure 4I). Together, these data suggest that CAR T-cell treatment triggers remodeling of the BM TME, associated with acquisition of widespread T-cell exhaustion, affecting both CAR T cells and endogenous T cells.

**Figure 4.**
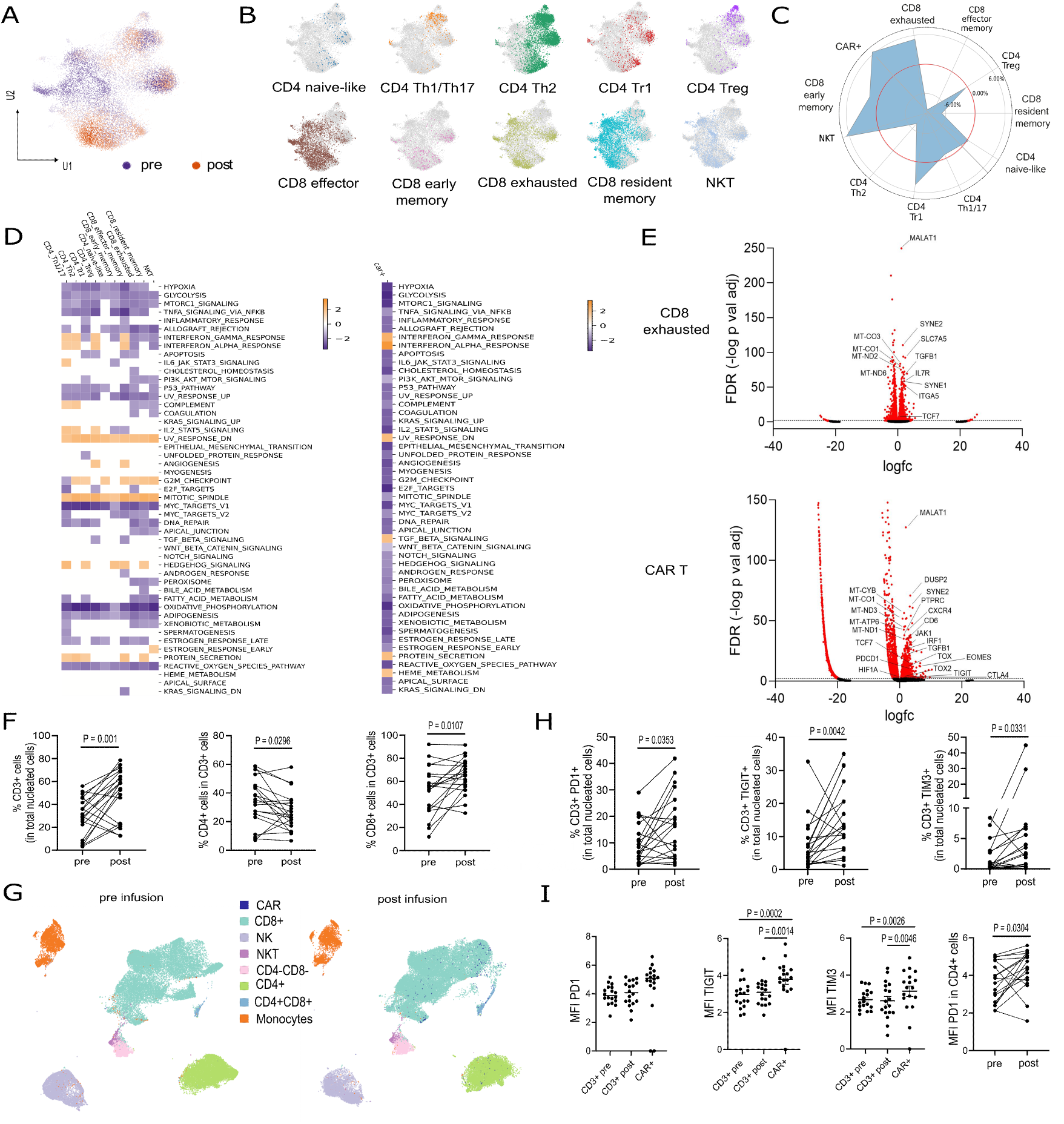
The TME post CAR T-cell treatment is characterized by Increased T-cell Exhaustion. (A) UMAP visualization of individual cells in the CD45+CD3+ libraries before and after anti-CD19 CAR T-cell treatment. (B) Garnett cluster-extended type classification of the CD3+ subpopulations. (C) Relative frequency of CD45+CD3+ cell subtypes. Delta proportion to estimate abundance reveals an increase in exhausted CD8+ T cells, CD4+ Type 1 regulatory T cells, and NKT after CAR T-cell treatment. (D) Significantly-enriched Molecular Signatures Database Hallmark gene sets in post-treatment versus pre-treatment CD45+CD3+ libraries within each immune cell types (left) and in CAR T cells (right). (E) Volcano Plots of differentially expressed genes (DEGs) in CD8 exhausted cell cluster (up) and CAR T cells (below) indicating genes up-regulated (rightward direction) and genes down-regulated (leftward direction) post CAR T-cell infusion compared to pre infusion. (F) Flow cytometry analysis (n=20 pre- and n=20 post- treatment paired BM samples) with Infinicyt depicting the frequencies of CD3+ cells in total nucleated cells and of CD4+ and CD8+ in CD3+ cells. Wilcoxon matched-pair t test. (G) UMAP representation with FlowSOM overlay of randomly selected cells (55000 per file) from 36 paired samples comparing before and after treatment. (H) Flow cytometry analysis with Infinicyt depicting frequencies of CD3+PD1+, CD3+TIGIT+, CD3+TIM3+ cells in total nucleated cells. Wilcoxon matched-pairs signed-rank test. (I) FlowSOM-generated MFI analysis of PD1, TIGIT and TIM3 in CD3+ cells before and after treatment and in CAR T cells. Statistical analysis was performed using one-way ANOVA with Friedman multiple comparison. MFI of PD1 in CD4+ cells before and after treatment. Statistical analysis performed using Wilcoxon matched-pairs signed-rank t test.

### PD1 expression on endogenous T cells is associated with lack of a durable response

We next evaluated if the remodeling of the microenvironment observed post-CAR T-cell infusion was associated with features of the TME pre-treatment. We found a strong correlation between the frequency of MDSCs, and similarly between the MFI of PD1, TIGIT, and TIM3 on endogenous T cells before and after treatment (SFigure 9A). An association was observed between the acquisition of TIGIT on resident T cells and on CAR T cells after infusion. The % and MFI of TIGIT in resident T cells correlated with that in CAR T cells, again suggesting an effect of the microenvironment in driving widespread T-cell exhaustion (SFigure 9B). PD1 and TIGIT MFI correlated with their % (SFigure 9C).

Given our hypothesis that the interplay between anti-CD19 CAR T cells and the immune compartment of the TME affects long-term therapeutic efficacy in B-ALL, we assessed the role of TME factors on the duration of response (DOR) and the event-free survival (EFS) in the cohort of patients analyzed by flow cytometry (STable 3). Specifically, we analyzed the frequency of MDSCs, CCR2+ monocytes, T cells and CAR T cells expressing PD1, and TIGIT post treatment. We found that a high % of CCR2 expression in monocytes after CAR T-cell treatment was only weakly associated with an increased DOR (hazard ratio (HR), 0.96, 95% CI, 0.92-1.00, p=0.05) but the association was stronger on EFS (HR, 0.95, 95% CI, 0.91-0.99, p=0.02). Conversely, higher % of CD3+PD1+ cells in endogenous T cells were significantly associated with decreased DOR (HR, 1.05, 95% CI, 1.01-1.10, p=0.03) and EFS after CAR T-cell treatment (HR, 1.05, 95% CI, 1.01-1.10, p=0.03, Table 1). The median survival was 1.13 years (0.24-3.46).

**Table 1:**
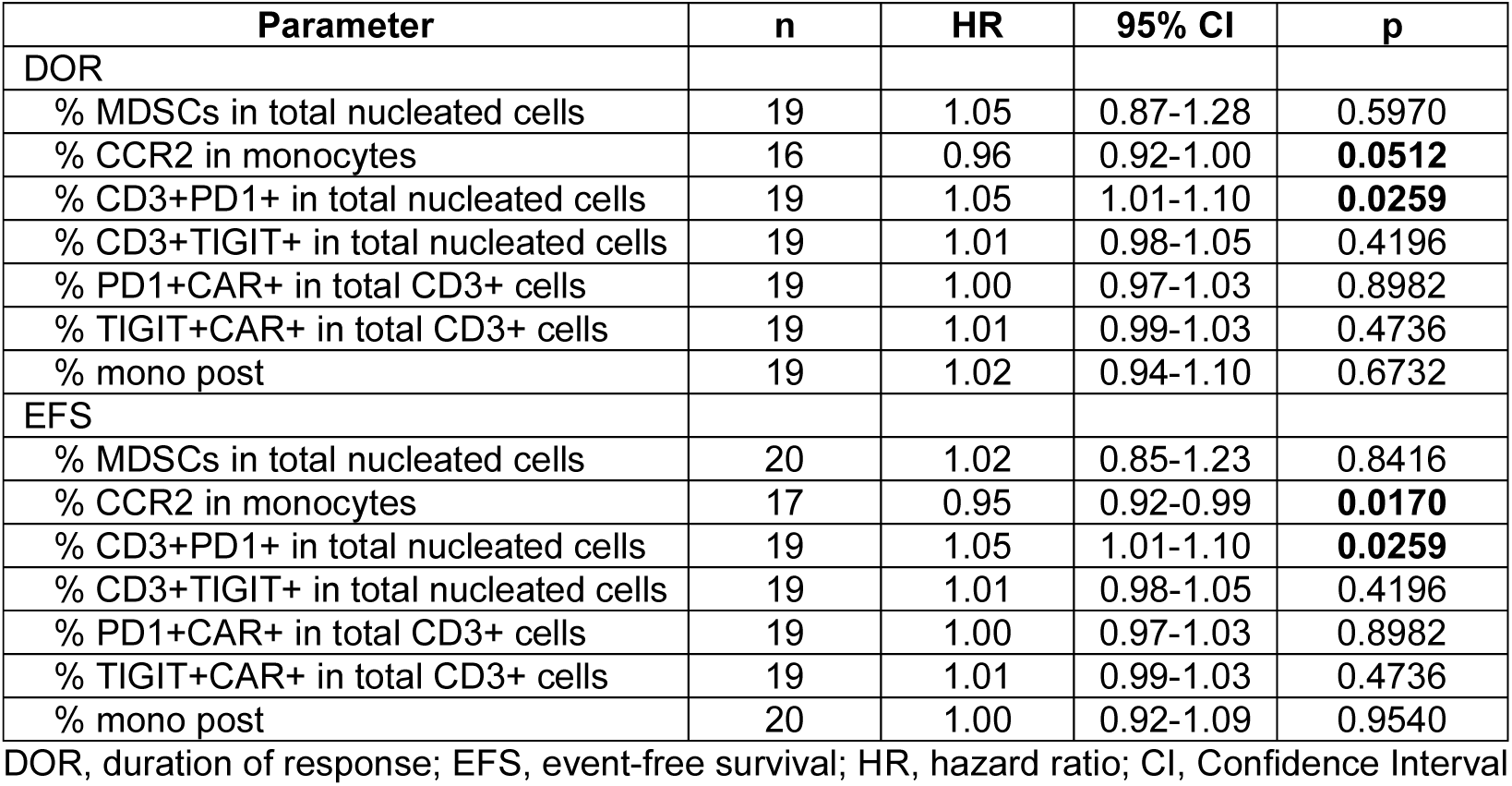
Results of the Cox model assessing the association between TME biomarkers analyzed with flow cytometry and DOR or EFS.

| Parameter | n | HR | 95% CI | p |
| --- | --- | --- | --- | --- |
| DOR |  |  |  |  |
| % MDSCs in total nucleated cells | 19 | 1.05 | 0.87-1.28 | 0.5970 |
| % CCR2 in monocytes | 16 | 0.96 | 0.92-1.00 | **0.0512** |
| % CD3+PD1+ in total nucleated cells | 19 | 1.05 | 1.01-1.10 | **0.0259** |
| % CD3+TIGIT+ in total nucleated cells | 19 | 1.01 | 0.98-1.05 | 0.4196 |
| % PD1+CAR+ in total CD3+ cells | 19 | 1.00 | 0.97-1.03 | 0.8982 |
| % TIGIT+CAR+ in total CD3+ cells | 19 | 1.01 | 0.99-1.03 | 0.4736 |
| % mono post | 19 | 1.02 | 0.94-1.10 | 0.6732 |
| EFS |  |  |  |  |
| % MDSCs in total nucleated cells | 20 | 1.02 | 0.85-1.23 | 0.8416 |
| % CCR2 in monocytes | 17 | 0.95 | 0.92-0.99 | **0.0170** |
| % CD3+PD1+ in total nucleated cells | 19 | 1.05 | 1.01-1.10 | **0.0259** |
| % CD3+TIGIT+ in total nucleated cells | 19 | 1.01 | 0.98-1.05 | 0.4196 |
| % PD1+CAR+ in total CD3+ cells | 19 | 1.00 | 0.97-1.03 | 0.8982 |
| % TIGIT+CAR+ in total CD3+ cells | 19 | 1.01 | 0.99-1.03 | 0.4736 |
| % mono post | 20 | 1.00 | 0.92-1.09 | 0.9540 |

DOR, duration of response; EFS, event-free survival; HR, hazard ratio; CI, Confidence Interval

### Modeling intercellular communication suggests induction of a chronically inflamed, hypoxic environment driving T-cell dysfunction

To unravel the mechanisms leading to the T-cell dysfunction associated with decreased long-term therapeutic efficacy, we explored the intercellular communication between different cell compartments, applying the computational method NicheNet to our scRNA-seq data.^27^ By using all cellular components of the niche as sender, and myeloid cells as receiver, we found CAR T cells and endogenous T cells to contribute to induction of *HIF1α*, *STAT1* and *IL6R* expression in myeloid cells (Figure 5A). Notably, TGF-β and IFN-γ expressed by T cells were predicted to be part of the network of communication signals mediating this effect (SFigure 10A). We then used NicheNet to predict signaling pathways from the niche influencing CAR T-cell fate and behavior. We observed induction of *NR4A2*, and *IRF1*, previously associated with T-cell exhaustion,^28,29^ *TGFB1*, the BM-attractant *CXCR4*, *SLC7A5* transmembrane amino acid transporters, and *ITGA4* (Figure 5B and SFigure 10B). *TGFB1*, *SLC7A5*, and *ITGA4* were also induced by the BM niche in endogenous exhausted T cells, in parallel to *TGFBR2*, and *NFKB* (SFigure 10C). Interestingly, myeloid cells were the main driver for the generation of an autocrine TGF-β signaling in both CAR T cells and endogenous T cells (SFigure 10B-D). Consistent with DEG analysis in the myeloid cluster, VEGFA, VEGFB and CCL2 were again identified as key players. We detected a slight increase of phospho-smad signal post-treatment (SFigure 10E), associated with Smad2 and Smad3 upregulation (SFigure 10F). The generalized increase in *HIF1α*, *TGFB1*, and *TGFBR2* expression (Figure 5C) was validated by digital droplet (dd)-PCR in purified CD3+ and CD3- BM cell subsets (SFigure11A).

**Figure 5.**
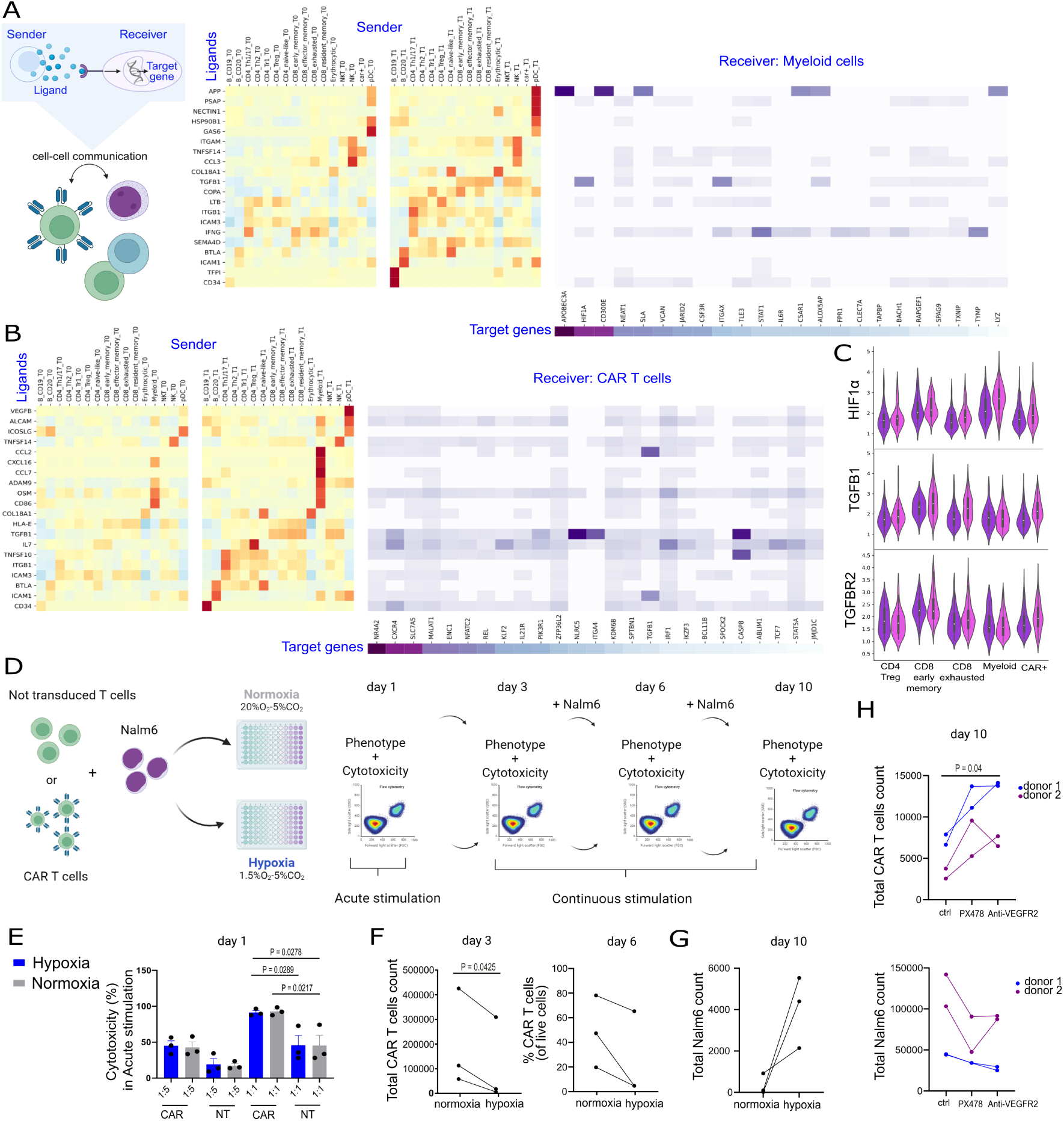
NicheNet-inferred cell-to-cell interactions revealed a role of chronic inflammation and hypoxia in driving T cell dysfunction. (A) NicheNet’s ligand activity prediction (left) and ligand-target matrix (right) assessed using all niche as sender and myeloid compartment as receiver and (B) using all niche as sender and CAR T cells as receivers. Heatmap visualization of prioritized ligands’ gene expression levels (Z-score) in specific cell compartments of all niche (left) and of predicted ligand activity in the receiver population (right). (C) Violin plots of scRNA-seq data depicting *HIF1α*, *TGFB1* and *TGFBR2* expression distribution in different immune cell clusters before and after treatment. (D) Schematic outline of the co-culture experiment simulating acute and chronic stimulation under normoxic and hypoxic conditions, in which CAR T cells are stimulated with Nalm6 cells for 24 hours (acute) or repeatedly every 3 days (chronic) at an effector:target (E:T) ratio of 1:1. (E) Cytotoxicity of CAR T cells after 24 hours of co-culture with Nalm6 cells at different effector to target (E:T) ratios compared with not-transduced (NT) cells (left). One-way ANOVA with Tukey multiple comparison. Data in E illustrate the mean ± SD of triplicates from three different donors. Paired t test. (F) Flow cytometry analysis depicting the CAR T-cell count and frequency after 3 and 6 days of co-culture, respectively. (G) Flow cytometry analysis depicting the outgrowing Nalm6 cell count after 10 days of co-culture. Data in F-G illustrate the mean of triplicates from three different donors. Paired t test. (H) Flow cytometry analysis depicting the CAR T-cell count and outgrowing Nalm6 cell count after 10 days of co-culture in the absence or presence of HIF1α inhibitor (PX478) or anti-VEGFR2 antibodies at an E:T ratio of 1:3. Data are presented as duplicates from two different donors. P value is determined by paired non parametric test (Friedman test with multiple comparison).

To determine whether hypoxia drives CAR T-cell exhaustion upon chronic T-cell activation, anti-CD19 CAR T cells were repeatedly stimulated with target cells, consisting of the B-ALL Nalm6 cell line, in normoxic (20% O_2_) and hypoxic conditions (1.5% O_2_, Figure 5D). Short-term co-culture in hypoxic conditions did not affect CAR T-cell functionality (Figure 5E). However, decreased proliferation was observed upon persistent antigen exposure under hypoxic conditions, resulting in decreased % of CAR T cells (Figure 5F and SFigure 11B), increased expression of TGFBR2 (SFigure 11C) and a higher proportion of terminal effector CD3+CAR+ cells (SFigure 11D). Further, we observed decreased CAR T-cell functionality associated with incomplete clearance of target cells at the end of the assay (Figure 5G). Therefore, we assessed whether inhibition of HIF1α or blockade of the VEGF-A/VEGFR2 interaction prevents T-cell dysfunction. CAR T-cell proliferation, together with target cell clearance, was increased by anti-VEGFR2 blocking antibodies ^30^ and, to some extent, by pharmacological inhibition of HIF1α with PX478 ^31^ during persistent antigen exposure under hypoxic conditions (Figure 5H). Thus, CAR T-cell mediated inflammation favors a hypoxic microenvironment that, during persistent stimulation, drives T cells towards a dysfunctional state.

### Tumor-bearing HSPC-humanized mice display increased monocytes and MDSCs following CAR T-cell therapy

To validate the direct effect of CAR T cells in inducing the observed TME remodeling in the absence of lymphodepletion and bridging therapy, we reconstituted newborn NSG mice by intrahepatic injection of CD34+ HSPCs and, after establishment of human hematopoiesis (SFigure 12A), transplanted with Nalm6, modified to express luciferase (Figure 6A). To avoid xenograft rejection, n-hu-Nalm-NSG mice received 3 x 10^6^ anti-CD19 CAR T cells generated from the splenocytes harvested from mice humanized with the same HSPC donor (SFigure 12B-C). CAR T cells mediated robust anti-leukemic activity in vivo, as demonstrated by the flux signal, compared to mice treated with NT cells (Figure 6B and SFigure 13A). At the time of euthanization, mice treated with CAR T cells showed almost complete clearance of leukemic blasts in the BM, but healthy B-cell reconstitution, associated with absence of human CD3+ T-cell engraftment (Figure 6C-D). Notably, we observed an increased frequency of human monocytes in the BM of CAR T cell-treated mice compared with control mice, associated with increased percentage of VEGFR2+CD33+ cells (median 5.2%, SD 2.8% in CAR T vs NT 0.9%, SD 0.6%) and HIF-1α^high^CD33+ cells (6.9 %, SD 0.5% vs 1.3%, SD 1.9 %), as well as MDSCs (0.8%, SD 0.5% vs 0.3%, SD 0%, Figure 6E and SFigure 13B-C). Purified myeloid cells were analyzed with bulk transcriptomics and confirmed our abovementioned scRNA-seq data, including increased expression of CCL2 and genes involved in biological oxidation and steroid biosynthesis (*CYP26B1, CYP27C1*), ROS (*SLAMF8, PXDNL, MAOA*), and MDSCs (*MMP19, NOS1AP*) (Figure 6F and SFigure 13D).

**Figure 6.**
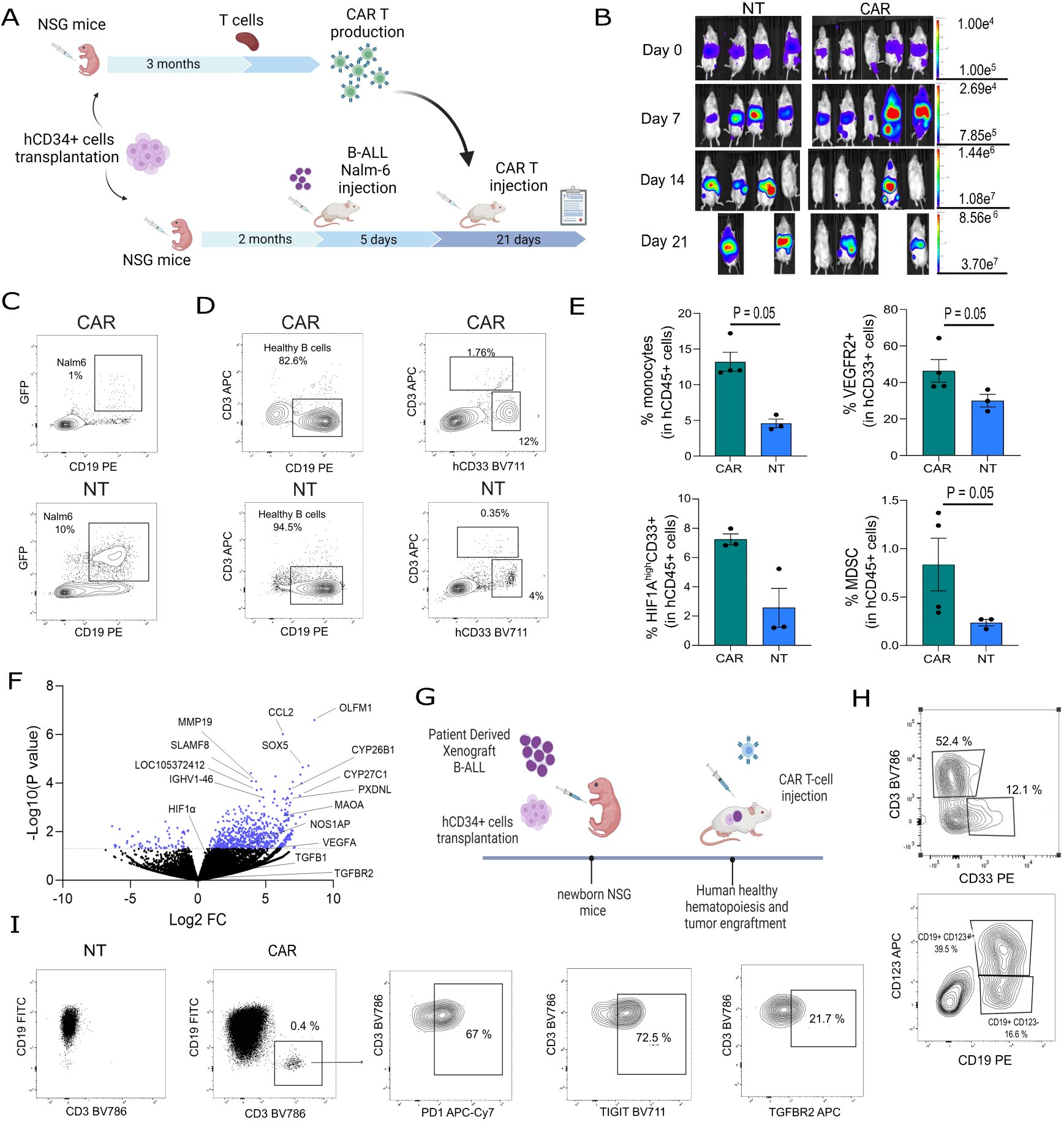
CAR T cells treatment remodeled the TME in tumor-bearing HSPC-humanized mice. (A) Newborn NSG mice were sub-lethally irradiated, injected intrahepatically with human PB-mobilized CD34+ cells and, after establishment of human hematopoiesis, infused with Nalm6 Luciferase+ cells. At day 5, mice were injected with 3×10^6^ CAR T cells or NT cells, generated from splenocytes harvested from mice humanized with the same HSPC donor by lentiviral transduction of an anti-CD19CAR.BBz. (B) IVIS images of mice injected with CAR T cells or NT cells. (C-D) Representative flow cytometry plots of GFP+CD19+ tumor cells, GFP-CD19+ healthy B cells, CD33+ myeloid and CD3+ T cells in BM of animals from each treatment at euthanization. One representative mouse per group is shown. (E) Percentage of CD33+HLA-DR+CD14+/CD15+ monocytes cells, VEGFR2+ in CD33+ myeloid cells, HIF1α^high^CD33+ cells and CD33+ HLA-DRlow/CD14+/CD15+ MDSC in BM of mice treated with CAR T or NT cells at euthanization. (F) Volcano plot of DEGs in myeloid cells purified from the BM of treated mice with CAR T cells compared with mice treated with NT cells. (G) Newborn NSG mice were sub-lethally irradiated, and injected intrahepatically with human PB-mobilized CD34+ cells and BM cells from secondary PDX recipient. After tumor engraftment, mice were injected with CAR T cells or NT cells generated from HSPC-humanized mice. (H) Representative flow cytometry plots of T cells, myeloid cells,CD123+CD19+ tumor cells, CD123-CD19+ B healthy cells in PB of a representative mouse at 2 months post transplantation. (I) Representative flow cytometry plots of CD3+ cells expressing PD1, TIGIT, TGFBR2 in BM of a representative mouse treated with CAR T cells at euthanization.

Since T cells were no longer detected after tumor clearance in the n-hu-Nalm-NSG model, we developed combined HSPC-humanized / PDX mice, by concomitant intrahepatic injection of CD34+ HSPCs and BM cells from secondary PDX recipient (Figure 6G). Remarkably, our n-hu-PDX-NSG model recapitulates the human tumor immune microenvironment, reconstituting a functional human immune system, including CD3+ T cells, CD19+ B cells and CD33+ myeloid cells, as well as the human CD19+CD123+ tumor (Figure 6H). In this model, at sacrifice all treated mice showed substantial leukemia infiltration in the BM with absence of other cell compartments, except for CD3+ cells in CAR T cell-treated mice. As seen in the patient samples, T cells showed high expression of TIGIT (median 79.4% of TIGIT+CD3+ cells, SD 13.4%), PD1 (47.1%, SD 18.1%), and TGFBR2 (55.8%, SD 17.1%, Figure 6I).

Overall, our mouse data provide experimental confirmation of the remodeling of the BM immune niche after CAR T-cell treatment, characterized by increased hypoxia, leading to skewing of the myeloid compartment towards MDSCs, and increased numbers of exhausted T cells.

## Discussion

In our study, single-cell transcriptomic and spectral flow-cytometry analyses uncovered previously unappreciated, extensive remodeling of the BM-resident immune niche in response to CAR T-cell mediated immune cell activation in B-ALL. We observed an increase in myeloid cells, particularly monocytes and MDSCs. GSEA on the myeloid cell subset indicates an inflammatory activation in response to type 1 and 2 IFNs, in line with recent studies,^32^ but also, hypoxia, which paradoxically results in a metabolic switch towards lipid metabolism, and conversion to MDSCs. The observed shift towards lipid metabolism through beta-oxidation is characteristic of tumor-infiltrating MDSCs and also modulates their immunosuppressive functions.^33^

Along with myeloid cells, we observed that type 1 and 2 IFN signaling results in activation of endogenous T-cell immunity, in line with our PB data, and an acquisition of features associated with T-cell exhaustion, including increased PD1 and TIGIT expression, widespread inhibition of glycolysis and OXPHOS,^22^ but preserved lipid metabolism, consistent with an adaptive response to energy deprivation.^6^ Similar but more pronounced changes were observed in CAR T cells, including upregulation of *TOX, HIF1α, EOMES* and exhaustion markers, in line with previous data in other malignancies indicating an evolution of CAR T cells toward a dysfunctional state in the TME.^34–36^ Pre-treatment systemic inflammation, IFN signaling and expansion of MDSC have recently been associated with reduced CAR T-cell expansion and lack of durable response in large B-cell lymphoma.^37^ Our data expand these findings by revealing generalized immunosuppression in the TME in response to CAR T-cell mediated bystander activation of endogenous immunity, as in chronic inflammation.^38,39^

The acquisition of exhaustion in endogenous T cells after CAR T-cell infusion appears to be clinically relevant in our cohort, as we find that an increased proportion of PD1+ T cells has a higher risk of shorter DOR and EFS. These data of ours present remarkable similarities with previous observations showing increased non-CAR CD8+ TCF1+ T cells and TCR diversity in myeloma patients treated with anti-BCMA CAR T cells.^40^ Moreover, a less exhausted phenotype in CD8+ T cells correlated with longer PFS, while systemic T-cell exhaustion was associated with poor CAR T-cell expansion in myeloma and B-cell lymphoma patients, respectively.^40–42^ Therefore, these findings suggest a role of induced endogenous immunity in influencing CAR T-cell activity, and highlight the importance of incorporating strategies that dampen excessive inflammation in the TME to prevent CAR T-cell exhaustion and promote beneficial rather than detrimental endogenous T-cell recruitment.^43–46^

By modeling intercellular communications, we noticed that HIF1α/VEGF up-regulation in myeloid cells and TGFB1/TGFBR2*-*autocrine signaling in T cells were interconnected and could lead to T-cell dysfunction. HIF1α, VEGF, and TGF-β have been identified as key drivers of MDSC differentiation and functions in the TME,^47,48^ and as factors promoting T-cell exhaustion.^49^ In inflamed tissues and cancer, increased oxygen consumption results in hypoxia, which, together with chronic antigen stimulation, alters immune cell function, trafficking, and metabolism, leading to exhaustion. Intriguingly, hypoxia, TGF-β, and myeloid inflammation pre-treatment were reported to be negatively associated with EFS and DOR in patients with large B-cell lymphoma treated with axi-cel.^50^ Our mechanistic in vitro studies suggest that increased hypoxia post-treatment drives the acquisition of a dysfunctional state in CAR T cells when combined with chronic stimulation. Importantly, inhibition of either VEGFR2 or HIF1α reinvigorated CAR T cells and restored their anti-tumor functions. These data are further corroborated by our results in tumor-bearing HSPC-humanized mice showing accumulation of VEGFR+ and HIF1α^high^ myeloid cells, MDSCs and acquired expression of PD1, TIGIT, and TGFBR2 on T cells after CAR T-cell infusion.

Whilst providing valuable new insight into the crosstalk between the BM microenvironment and CAR T-cells in B-ALL, our study has limitations: it has been conducted in a heterogeneous cohort including a limited number of patients and is therefore restricted to describing associations. Hence, validation in additional cohorts and analysis of similar datasets would strengthen the relevance of our findings and to draw causal conclusions.

Overall, our data suggest that CAR T cells promote myeloid, and MDSC expansion in the BM microenvironment in patients with B-ALL, ultimately leading to T-cell exhaustion as an active feedback loop in response to CAR T-cell-mediated inflammation. Mechanistically, the acquisition of an exhausted phenotype in endogenous T cells and CAR T cells is induced by the coordination of chronic inflammation and microenvironmental stressors such as HIF1α, VEGF, and TGF-β and signaling, and may antagonize the effect of the therapy.

## Acknowledgments

We are grateful to the patients and patients’ families.

This work was supported by AIRC/Cancer Research UK (CRUK)/Spanish Association Against Cancer Scientific Foundation (FC AECC), grant number 22791 (Accelerator award) to AB and CFM, Ministero della Salute GR-(2016-02363491) to CFM, Swiss National Science Foundation (PR00P3_201621) to CFM, the Helmut Horten Foundation to CFM, Fondazione M. Tettamanti De Marchi ONLUS. M. Ponzo was supported in parts by funds from EHA (Research Mobility Grant), and AIRC (Fellowships for Abroad Post-Doc 2023).

## Author contributions

M. Ponzo designed and performed all the experiments, collected and analyzed the data and wrote the manuscript. L.D. analyzed the scRNA-seq data, and wrote the manuscript. C. Buracchi assisted with spectral flow cytometry. FlowSOM analysis was carried out by M.M.S. and S.N., who also analyzed bulk RNA-seq data. C. Bugarin assisted with phospho-flow analysis. R. Bason assisted with scRNA-seq library generation. G. Rossetti and R. Bonnal assisted with the scRNA-seq data analysis. B.R. and A.M. provided information on patient’s clinical data. C.P. assisted with animal experiments. M.G.M. wrote and approved the manuscript. G. Risca and S.G. performed statistical analysis. A.B. and A.R. conceptualized the clinical studies. A.B. and G.G. conceptualized the study and obtained financial support. M. Pagani conceptualized the study and led the scRNA-seq data collection, analysis and interpretation. C.F.M. conceptualized the study, designed the experiments, analyzed the data, obtained financial support, wrote and approved the manuscript. All authors reviewed and approved the final manuscript.

## Disclosure of Conflicts of Interest

AR has received honoraria for lectures from Novartis, Bristol Myers Squibb, Sanofi, Abbvie, Amgen, Pfizer, Kite-Gilead, Jazz, Astellas, Incyte, and Omeros. M Pagani is of the board of Directors and stakeholder of CheckmAb s.r.l. and is a recipient of a grant under a research agreement with Bristol-Myers Squibb Company. AB serves in the scientific advisory board (SAB) of CoImmune and Galapagos. CFM, MP, GG and AB reports a patent pending for methods to predict and improve the efficacy of CAR T-cell therapy by combining inhibitors of hypoxia, VEGF pathways, and inflammation in the tumor microenvironment. CFM and AB are inventors on a patent related to non-viral CAR-T cell therapy (European patent application 15801344), filed by the Tettamanti Foundation. The remaining authors declare no potential conflicts of interest.

## Data Sharing Statement

Data generated or analyzed during this study are included in the article and Supplementary Material. The scRNA-seq data generated in this study and code used for analysis are available into the ArrayExpress collection in BioStudies with the accession number “E-MTAB-14549” (https://www.ebi.ac.uk/biostudies/arrayexpress/studies/E-MTAB-14549).

## Notes

https://www.ebi.ac.uk/biostudies/arrayexpress/studies/E-MTAB-14549

