## Supplementary data for "Acquisition of an immunosuppressive microenvironment after CAR-T therapy drives T-cell dysfunction and resistance"

Marianna Ponzo et al.

Corresponding authors:

Chiara F Magnani,

Andrea Biondi,

The file includes

Methods

Supplemental Table 1 to 8

Supplemental Figure 1 to 13

**Methods:**

**Patient’s Peripheral blood** **analysis**

All patients or their legal guardians provided written informed consent for sample collection and analysis. Studies were in accordance with the declaration of Helsinki and Ethical approval was obtained. Patients underwent CAR T treatment at Fondazione Monza e Brianza per il Bambino e la sua Mamma (MBBM), Monza, Italy (pediatric patients) or at Azienda Socio Sanitaria Territoriale Papa Giovanni XXIII, Bergamo, Italy (adult patients). Peripheral blood (PB) of B-ALL patients who received CARCIK-CD19 or autologous CAR T cells was used to analyze the kinetics over time of total WBC, lymphocytes, monocytes, and neutrophils. CAR T and CD3+ kinetics were assessed by flow cytometry. Hematology laboratory data during the first two months after CAR T-cell infusion were routinely monitored for clinical purposes. Cell counts were performed with Beckman Coulter Uni Cel DxH 800.

**Frozen BM Preparation, Flow Cytometry, and Cell Sorting**

For scRNA-seq analysis, samples were selected according to the presence of detectable CAR T cells and CD45+CD3- in the BM sample post-infusion. Material consisted of frozen BM samples. Cell counts and viability after sorting were optimal and sufficient to proceed with the scRNA-seq analysis. Frozen human BM and IP cryotubes were rapidly thawed and stained with hCD45 PO (clone HI30, Invitrogen) and CD3 PerCP (clone SK7, BD) or with CD19 biotin protein (Miltenyi) followed by Biotin APC (REA746, Miltenyi) and CD3 PercP (clone SK7, BD) antibodies, respectively. BM cells were separated into CD45+CD3+ and CD45+CD3- fractions, whereas IP was separated into CD3+CAR+ and CD3+CAR- cells by flow cytometry-based sorting (BD FACS ARIA III analyser). All samples were gated based on physical parameters, followed by the exclusion of doublets and death cells (DAPI high). After sorting, cells were counted and checked for viability.

**scRNA-seq Data Processing and Quality Control**

To enable identification of cells expressing the anti-CD19 CAR transcripts, sequences corresponding to the recombinant mouse CDR3 region directed against CD19 molecule and the leader were added to the reference genome and annotated. Individual libraries were then analyzed using the Scanpy package 9 and putative doublets were removed using the scrublet package with the threshold set to 0.2 10,11. To limit technical artifacts, libraries were then downsampled to have average counts per cell values comparable to the least represented library (~2500 counts per cell on average). All libraries were subsequently concatenated into a single dataset. Genes expressed in less than 1/1000 cells (i.e., 72 cells) were removed as low expressed genes. To exclude low quality cells, barcodes with less than 200 genes were removed. Furthermore, cells deviating for more than a fixed number of mean absolute difference (MAD) from the median of selected metrics were excluded from the analysis. The metrics evaluated percentage of mitochondrial transcripts (nMAD=4, threshold ~ 20%), percentage of ribosomal transcripts (nMAD=2, threshold ~ 50%), number of UMI (nMAD=3, threshold ~ 6000), number of expressed genes (nMAD=3, threshold ~ 2700). In the end 71,407 cells were retained for further analysis. The data were normalized to a total expression of 10^4^ counts per cell and log transformed. Highly variable genes (HVG) were selected using the corresponding function from the scanpy package, increasing the min_mean parameter to 0.1 which yielded 909 HVG. For visualizing the data, the dimensionality of the matrix was reduced further to project the cells in two-dimensional space using PCA followed by UMAP (considering 25 PCs, 30 nearest neighbours, min_dist=0.1). To reduce the effect of technical artifacts onto cell embeddings, the HARMONY algorithm12 was applied to the dataset before UMAP projection to achieve better integration.

**Cell type annotation**

For the CD3- compartment, clusters were assigned to identified cell types based on the expression of defined population markers using unsupervised clustering ^1^. For the classification of cells from the CD45+/CD3+ compartments, we relied on a previously described strategy employing a hierarchical logistic classifier^2^. Briefly, at each level of the cellular hierarchy, a set of positive and negative markers were selected to identify populations of interest (marker file) and were used to select high-confidence cells from a reference dataset (GSE161529, Supplemental Table S4). Residual cells not assigned to any of the target classes were further analyzed and manually annotated according to differentially expressed gene markers. For MDSC classification, the signature transcriptional activity was evaluated at single cell level by taking the singular value decomposition (SVD) score on the signature-restricted gene expression matrix.

**Differential abundance evaluation**

To evaluate differential cell-type abundance across time points in CD3- cells, libraries of interest were subset and retained from the full dataset and both PCA and UMAP representation were computed. MILO algorithm was then employed to highlight cellular populations with differential representation across conditions (design= ~0+timepoint, contrast= timepointT1-timepointT0). To identify neighborhoods we considered k= 30 nearest neighbours over 30 PCs and set the attribute prop= 0.1 while all other parameters were left as default.

Due to the higher transcriptional homogeneity observed in CD3+ data subset, we evaluated differential abundance considering the overall relative proportion of a specific cell subtype across time points.

**Cellular crosstalk evaluation**

To predict intracellular communication, we used NicheNet, a computational tool for predicting ligand-receptor binding and target genes among interacting cells by combining their expression data with prior knowledge of signaling pathways and gene regulatory networks ^2^. Of interest, the recently added ‘MultiNicheNet’ procedure allows to compare signaling within a specific niche across different condition (time points in our case). We defined different niches in which, for each iteration, we focused our analysis on a specific cell type as receiver (myeloid, CD8_exhausted, and CAR T cells respectively) while leaving all other cell types as possible senders. This approach highlights ligand-receptor pairs, which showed differential expression across conditions and were likely to induce the expression of observed DEG target for the receiver population. MultiNicheNet analysis was performed setting expression_pct = 0.10, specificity_score_LR_pairs = "min_lfc", include_spatial_info_sender and include_spatial_info_receiver both set to FALSE, lfc_cutoff = 0.15, specificity_score_targets = "min_lfc", top_n_target = 250. Relative weights for prioritization were set at scaled_ligand_score= 4, scaled_ligand_expression_scaled= 1, ligand_fraction= 1, scaled_ligand_score_spatial= 0, scaled_receptor_score= 0.5, scaled_receptor_expression_scaled= 0.5, receptor_fraction= 1, ligand_scaled_receptor_expression_fraction= 1, scaled_receptor_score_spatial= 0, scaled_activity= 0, scaled_activity_normalized= 1, bona_fide= 1

**Phenotypical Analysis**

To identify molecular pathways altered within cell types at different time points, we performed a GSEA analysis on DEG for each cell type (as returned by calling rank_genes_groups from the scanpy package onto a specific cell type grouped by time points). To evaluate different aspects of cellular behavior, we tested the Hallmark collection from the Molecular Signatures Database (MSigDB 15) by GSEA as well as the hand-refined and metabolically scoped KEGG and Reactome collections described in scMetabolism ^3^.

**CAR T-cell detection**

An in silico limiting dilution assay was performed to test the detection of CAR+ cells in a complex cellular mixture by diluting barcodes from the CD3+CAR+ IP with BM-derived CD45+ CD3+ and to evaluate the ability of unsupervised clustering techniques to efficiently partition CAR+ cells together.

For the identification of CAR+ cells in the final dataset, BM CD45+CD3+ libraries were preliminarily explored individually. Cells expressing the annotated CAR sequences were directly tagged as ‘cart’ cell. Following data clustering using unsupervised community detection algorithms (leiden algorithm 17), the ‘cart’ label was then extended to clusters enriched in CAR-expressing cells.

**Unsupervised flow-cytometric analysis with FlowSOM**

All fluorescence conjugated antibodies are listed in Supplemental Table S5 and Supplemental Table S6. We used the raw fcs files from 36 paired samples (pre and +1 month) from 18 patients as described above. First, we manually gated on live cells, according to physical parameters and CD45 expression, and applied a spillover matrix using polygonGate and compensation functions of flowCore package (v.2.12.0). We performed logicle transformation employing FCSTransTransform function. The quality of transformed fcs files was assessed using PeacoQC function of PeacoQC package (v 1.11.0). Processed fcs were aggregated together in a 2x10^6^ events file (roughly 55000 live cells per file). Clustering analysis was performed with FlowSOM R package (v2.8.0). The first clustering step was performed using a 15x15 grid producing 225 clusters and 10 metaclusters to identify Monocytes and Lymphocytes and to eliminate the remaining debris; we employed the same markers used in the manual gating strategy. A second level of clustering was performed on Monocytes (6x6 grid and 15 metaclusters) and Lymphocytes (15x15 grid and 15 metaclusters) to dissect their composition. For each level, counts, percentages and MFI were extracted from each subset using GetFeatures function of FlowSOM package. Downstream statistical analysis was performed using GraphPad Prism (version 10). UMAP was computed using the umap function from the uwot R package (v0.1.16.9) and the n_neighbors parameter set to 15.

**Analysis of TGF-β signaling**

Activation of the TGF-β signaling was verified through phospho-flow analysis of the Smad proteins (basal phosphorylation). To evaluate the phosphorylation of SMAD proteins, thawed BM cells were rested in RPMI medium containing 0.1% FBS for 2 hours at 37°C. Cells were then fixed with 1.5% paraformaldehyde and permeabilized with 90% ice-cold methanol prior to staining with anti-Smad2 (pS465/pS467)/Smad3 (pS423/pS425) antibody (clone O72-670,BD Biosciences) or isotype matched IgG, anti-CD7 ECD (Beckman Coulter) and anti-CD45 PerCP (BD).

**ddPCR analysis**

BM cells were separated into CD45+CD3+ and CD45+CD3- fractions through CD3+ positive selection on LS column with CD3 MicroBeads (Miltenyi). RNA was extracted with miRNeasy micro kit (QIAGEN). 10 ng of RNA was retrotranscribed with SuperScript™ IV Reverse Transcriptase (Thermo Fischer) and Digital Droplet ™ PCR (ddPCR™) (BIO RAD) was performed for the evaluation of HIF1α (dHsaCPE5033624), TGFB1 (dHsaCPE5055188), VEGFA (dHsaCPE5034754) genes. Analyses were performed with QuantaSoft Analysis pro Software ™ (BIORAD).

**In vitro Co-culture**

anti-CD19 CAR T cells were manufacture as previously described in ^4,5^. T cells were obtained from anonymized healthy donors from the blood donation service. CAR T cells were cultured with Nalm-6 for 24 hours (acute stimulation) at effector to target (E:T) ratios of 1:1 and 1:5 under normoxia (20% O_2_) or hypoxia (1.5% O_2_) compared to NT cells. Cytotoxicity was calculated as 100-(% live Nalm6 in co-culture with effector cells /% live Nalm6 alone) ×100. To model persistent antigen exposure, CAR T cells were cultured with Nalm6 at a E:T ratio of 1:1 and restimulated with target cells at day 3 and day 6. All conditions were run in triplicates. For the drug treatment, CAR T cells were cultured with Nalm-6 at a E:T ratio of 1:3 under hypoxia (1% O2) in the presence of anti-VEGFR2 antibody (ramucirumab, Selleck Chemicals) or PX478 (CAT #S7612, Selleck Chemicals) at 10µM and 50 µg/ml, respectively, and re-stimulated with target cells at day 3 and day 6. All conditions were run with two donors in duplicates and anti-VEGFR2 antibody and PX478 were replenished at the same concentration every 36 hours or every re-stimulation, respectively.

Cells were stained with a 18-colors panel. All fluorescence conjugated antibodies are listed in Supplemental Table S7. Data were acquired on a Cytek Aurora (Cytek® Biosciences) and flow cytometric analyses were performed using Infinicyt (Cytognos) and FlowJo.

**Generation of CAR T cells from hematopoietic stem/progenitor cell (HSPC)-humanized mice**

Animal experiments were conducted in compliance with procedures approved by the Veterinäramt des Kantons Zürich, Switzerland (194/2018 and 121/2022) and by the Italian Ministry of Health (approval no. 907/2016-PR). The animals were kept in the animal facilities of the University Hospital of Zürich and the Schlieren Campus of Zürich and of the Milano-Bicocca University. Splenocytes were harvested from mice humanized with CD34+ HSPCs from residual, left-over mobilized peripheral blood cells collected from stem cell infusion bags and processed by the biobank of the department of Medical Oncology and Hematology, University Hospital Zurich. The donor gave written consent. Studies were in accordance with the declaration of Helsinki and approved by the Cantonal Ethical Board Zurich, Switzerland (2019-01744). Anti-CD19 CAR T cells were produced by transduction with lentiviral vectors as previously described ^5^. The viral vector encoding the anti-CD19 CAR (clone FMC63), including the 41BB costimulatory domain, and the RQR8 sequences previously cloned in the pCDH-EF1α-MCS-T2A-GFP lentiviral plasmid was produced as previously reported ^5^. T-cell activation was performed with CD3/CD28 Dynabeads® (Thermo Fisher Scientific) followed by transduction with LV in the presence of 8 µg/ml Polybrene (Santa Cruz Biotechnology, #134220) the following day. On day 3, beads were removed with magnets. LV CAR T cells were cultured for 7 days, purified with FITC-Micro Beads (Miltenyi) after incubation with hCD34 FITC (ThermoScientific) and purified cells were expanded for additional 5-7 days before cryopreservation or immediate use.

**Tumor-bearing HSPC-humanized mice**

For the experiment depicted in Figure 6a, newborn pups were sub-lethally irradiated with 150 cGy with an RS-2000 irradiator (Rad Source, Buford, GA, USA) and transplanted by intrahepatic injection of 650.000-1x10^6^ CD34+ HSPCs from mobilized peripheral blood cells collected as described above. Levels of chimerism were monitored by flow cytometry of PB post-trans plantation and when the level demonstrated successful engraftment (i.e. >1%human CD45 positive cells in the peripheral blood), mice were infused with 0.5x10^6^ luciferase-GFP expressing Nalm-6. Animals with positive luciferase signal on day 5 were allocated to receive 3x10^6^ purified CAR+ T cells generated from splenocytes collected from HPCS-humanized mice with the same HSPC donor. For weekly bioluminescent imaging (BLI), mice were anesthetized and received 150 mg/kg bodyweight D-luciferin (PerkinElmer, Inc, Wal tham, MA, USA) as intraperitoneal injection. Image acquisition was performed on a Xenogen IVIS 200 machine (PerkinElmer) with the Living Image 149 Software. Animals were euthanized 21 days after T cell transfer. Hematological organs (PB, BM, and spleen) were collected and processed for flow cytometry analysis. The PDX model was established with BM material (Patient ID #003) as previously described ^6^. For the experiment reported in Figure 6g, newborn pups were sub-lethally irradiated with 150 cGy and co-transplanted by intrahepatic injection of 750.000 CD34+ HSPCs and 750.000 BM cells from the secondary leukemic recipients. After tumor engraftment, mice were treated with 3x10^6^ anti-CD19 CAR T cells generated from splenocytes from mice humanized with the same HSPC donor. Animals were euthanized 24 days after T-cell transfer. For flow cytometry analysis, cells from BM were stained with fluorescence conjugated antibodies listed in Supplemental Table S8.

HIF1α was detected in BM purified cells by intracytoplasmic staining (Fixation/Permeabilization Solution Kit, BD Bioscience) with anti-hHIF1α PE (clone546-16, Biolegend).

**RNA-seq**

BM cells were thawed, negatively purified using anti PE-Micro Beads (Miltenyi) after incubation with anti-hCD19 and anti-mCD45 PE antibodies, and lysed directly with Qiazol (Qiagen). Total RNA was isolated and purified with miRNeasy micro Kit (Qiagen, Germany). For library preparation, we used Universal Plus Total RNA-Seq with NuQuant (Tecan Genomics). Analysis of RNA-seq data was performed using Galaxay Europe (<https://doi.org/10.1093/nar.gkae410>). Briefly, after quality check, transcripts were aligned using Bowtie2 (version 2.5.3). Subsequently, counts were extracted using FeatureCounts (Version 2.0.3) and differentially expressed genes were identified using Deseq2 (version 1.40.2). Gene set enrichment analysis was performed using Web based gene set analysis toolkit (<https://www.webgestalt.org/>).

**Statistics**

Statistical analyses were performed in Graph Pad Prism Version 10 (GraphPad Software, Inc., La Jolla, CA) and SAS (v 9.4). Quantitative variables were described in terms of means ± stardard deviation (SD) and the degree of linear association was quantified by the Pearson correlation coefficient. Between groups comparisons were performed using the t-test when the groups were two and the ANOVA for three groups. Paired comparisons were performed by means of the Wilcoxon paired test. EFS was defined as the time from the date of CAR T-cell infusion to the earliest of the following events: no Complete Remission (CR), relapse or death from any cause, while DOR started from CR to relapse or death due to any cause, whichever occurred first. Patients were censored at the last follow-up in case no events occurred. To assess the impact of continuous variables on DOR and EFS we used the Cox model.

**Supplemental Tables**

| Patient ID | Age (years) | Sex | Genetic abnormality | Clinical Study - Infusion product | Response | Analyses |
| --- | --- | --- | --- | --- | --- | --- |
| #001 | 2 | F | t(4,11) | FT01 CARCIK-CD19 | CR | Hematology and scRNAseq |
| #002 | 17 | F | Negative for all translocations | Commercial anti-CD19 CAR T | CR | Hematology and scRNAseq |
| #003 | 6 | F | Negative for all translocations | Commercial anti-CD19 CAR T | CR | Hematology and scRNAseq |
| #004 | 5 | F | t(9;18) | FT01 CARCIK-CD19 | NR | Hematology |
| #005 | 27 | M | hyperdiploid | FT01 CARCIK-CD19 | NR | Hematology |
| #006 | 56 | M | Negative for all translocations | FT01 CARCIK-CD19 | NR | Hematology |
| #007 | 62 | F | t(9;22) Bcr/Abl1 | FT01 CARCIK-CD19 | CR | Hematology |
| #008 | 45 | F | t(9;22) Bcr/Abl1 | FT01 CARCIK-CD19 | CR | Hematology |
| #009 | 10 | M | t(4;11) | FT01 CARCIK-CD19 | NR | Hematology |
| #010 | 7 | M | Complex karyotype | FT01 CARCIK-CD19 | CR | Hematology |
| #011 | 32 | F | Negative for all translocations | FT01 CARCIK-CD19 | NR | Hematology |
| #012 | 39 | F | Negative for all translocations | FT01 CARCIK-CD19 | CR | Hematology |
| #013 | 63 | M | t(9;22) Bcr/Abl1 | FT01 CARCIK-CD19 | CR | Hematology |
| #014 | 52 | F | Hyperdiploid | FT01 CARCIK-CD19 | CR | Hematology |
| #015 | 30 | M | Complex karyotype | FT01 CARCIK-CD19 | CR | Hematology |
| #016 | 28 | M | t(9;22) Bcr/Abl1 | FT01 CARCIK-CD19 | NR | Hematology |
| #017 | 35 | F | Not available | FT01 CARCIK-CD19 | NR | Hematology |
| #018 | 60 | F | t(9;22) Bcr/Abl1 | FT01 CARCIK-CD19 | CR | Hematology |
| #019 | 46 | M | t(9;22) Bcr/Abl1, p190, mut T315 I | FT01 CARCIK-CD19 | NR | Hematology |
| #020 | 36 | M | t(9;22) Bcr/Abl1 | FT01 CARCIK-CD19 | CR | Hematology |
| #021 | 37 | M | t(9;22) Bcr/Abl1 | FT01 CARCIK-CD19 | CR | Hematology |
| #022 | 38 | F | t(9;22) Bcr/Abl1 | FT01 CARCIK-CD19 | CR | Hematology |
| #023 | 26 | F | Negative for all translocations | FT01 CARCIK-CD19 | CR | Hematology |
| #024 | 4 | M | Ph-like | Commercial anti-CD19 CAR T | CR | Hematology |
| #025 | 13 | F | t(1;19) | Commercial anti-CD19 CAR T | CR | Hematology |
| #026 | 21 | M | Negative for all translocations | Commercial anti-CD19 CAR T | PR | Hematology |
| #027 | 0 | F | t(4;11) | Phase 1/2 anti-CD19 CAR T | CR | Hematology |
| #028* | 7 | M | Complex karyotype | Commercial anti-CD19 CAR T | CR | Hematology |
| #029 | 2 | M | t(11;19) | Commercial anti-CD19 CAR T | CR | Hematology |
| #030 | 1 | M | Negative for all translocations | Phase 1/2 anti-CD19 CAR T | CR | Hematology and flow cytometry |
| #031* | 4 | M | t(11;19) | Phase 1/2 anti-CD19 CAR T | CR | Hematology and flow cytometry |
| #032 | 8 | M | t(9;22) Bcr/Abl1 | Commercial anti-CD19 CAR T | CR | Hematology and flow cytometry |
| #033 | 10 | M | t(12;21) | Commercial anti-CD19 CAR T | CR | Hematology and flow cytometry |
| #034 | 4 | F | t(4;11) | Commercial anti-CD19 CAR T | CR | Hematology and flow cytometry |
| #035 | 8 | M | Negative for all translocations | Commercial anti-CD19 CAR T | NR | Hematology and flow cytometry |
| #036 | 1 | F | KMT2A rearrangement | Commercial anti-CD19 CAR T | CR | flow cytometry |
| #037 | 19 | F | Negative for all translocations | Commercial anti-CD19 CAR T | CR | flow cytometry |
| #038 | 11 | F | t(9;22) Bcr/Abl1 | Commercial anti-CD19 CAR T | CR | flow cytometry |
| #039 | 4 | M | t(9;22) Bcr/Abl1 | Commercial anti-CD19 CAR T | CR | flow cytometry |
| #040* | 8 | M | Complex karyotype | FT03 CARCIK-CD19 | CR | flow cytometry |
| #041 | 16 | F | Negative for all translocations | FT03 CARCIK-CD19 | CR | flow cytometry |
| #042 | 47 | F | t(9;22) Bcr/Abl1 | FT03 CARCIK-CD19 | CR | flow cytometry |
| #043 | 45 | F | t(9;22) Bcr/Abl1 | FT03 CARCIK-CD19 | CR | flow cytometry |
| #044 | 66 | M | Negative for all translocations | FT03 CARCIK-CD19 | CR | flow cytometry |
| #045 | 48 | M | Ph-; trisomy 8 monosomy 7 | FT03 CARCIK-CD19 | CR | flow cytometry |
| #046 | 42 | F | Negative for all translocations | FT03 CARCIK-CD19 | CR | flow cytometry |
| #047 | 58 | M | J beta 1.1 | FT03 CARCIK-CD19 | CR | flow cytometry |
| #048 | 59 | M | t(9;22) Bcr/Abl1 | FT03 CARCIK-CD19 | CR | flow cytometry |
| #049 | 49 | F | Mixed phenotype | FT03 CARCIK-CD19 | CR | flow cytometry |

Abbreviations: F, female; M, male; CR, complete response; NR, no response; scRNAseq, single-cell RNA sequencing

*patient #028 received commercial CAR T cells when he was 2 years old, but relapsed post treatment and was enrolled to CARCIK-CD19 (indicated here as patient #040)

**Supplemental Table 1:** Patients’ characteristics

| Patient ID | Sample type | Estimated Number of cells | Number of reads (X10^6^) | Total Genes Detected | Saturation | Sequencing  Q30 | Mapping Quality |
| --- | --- | --- | --- | --- | --- | --- | --- |
| #001 | IP CD3+ CAR+ | 11105 | 177 | 18061 | 64% | **V** | **V** |
| #001 | IP CD3+ CAR- | 7372 | 313 | 18428 | 80% | **V** | **V** |
| #001 | BM pre CD45+/low CD3- | 4913 | 161 | 18876 | 78% | **V** | **V** |
| #001 | BM pre CD45+ CD3+ | 5451 | 112 | 17685 | 70% | **V** | **V** |
| #001 | BM post CD45+/low CD3- | 5526 | 342 | 19928 | 63% | **V** | **V** |
| #001 | BM post CD45+ CD3+ | 6812 | 458 | 18997 | 82% | **V** | **V** |
| #002 | IP CD3+ CAR+ | 9132 | 386 | 20594 | 58% | **V** | **V** |
| #002 | IP CD3+ CAR- | 10497 | 444 | 21029 | 46% | **V** | **V** |
| #002 | BM pre CD45+/low CD3- | 6631 | 322 | 19084 | 82% | **V** | **V** |
| #002 | BM pre CD45+ CD3+ | 7048 | 325 | 19759 | 63% | **V** | **V** |
| #002 | BM post CD45+/low CD3- | 1071 | 404 | 17676 | 94% | **V** | **V** |
| #002 | BM post CD45+ CD3+ | 5222 | 314 | 18623 | 76% | **V** | **V** |
| #003 | IP CD3+ CAR+ | 9096 | 467 | 21371 | 63% | **V** | **V** |
| #003 | IP CD3+ CAR- | 10116 | 508 | 20993 | 76% | **V** | **V** |
| #003 | BM pre CD45+/low CD3- | 9002 | 462 | 21527 | 81% | **V** | **V** |
| #003 | BM pre CD45+ CD3+ | 10052 | 497 | 22346 | 62% | **V** | **V** |
| #003 | BM post CD45+/low CD3- | 5982 | 477 | 20084 | 89% | **V** | **V** |
| #003 | BM post CD45+ CD3+ | 7877 | 511 | 22313 | 71% | **V** | **V** |

Abbreviations: IP, infusion product; BM, bone marrow

**Supplemental Table 2:** Quality control data

| **Patient ID** | **DOR (years)** | **Event** | **EFS (years)** | **Event** |
| --- | --- | --- | --- | --- |
| #035 | . | . | 0.08 | 1 |
| #040 | 0.07 | 1 | 0.15 | 1 |
| #047 | 0.18 | 1 | 0.27 | 1 |
| #044 | 0.23 | 1 | 0.31 | 1 |
| #049 | 0.24 | 1 | 0.31 | 1 |
| #039 | 0.27 | 0 | 0.34 | 0 |
| #043 | 0.33 | 1 | 0.41 | 1 |
| #033 | 0.45 | 1 | 0.52 | 1 |
| #041 | 0.48 | 1 | 0.56 | 1 |
| #037 | 0.49 | 1 | 0.57 | 1 |
| #048 | 0.51 | 0 | 0.59 | 0 |
| #038 | 0.51 | 0 | 0.58 | 0 |
| #046 | 0.67 | 1 | 0.72 | 1 |
| #036 | 0.71 | 0 | 0.79 | 0 |
| #031 | 0.81 | 1 | 0.89 | 1 |
| #045 | 1.05 | 0 | 1.13 | 0 |
| #032 | 1.13 | 1 | 1.21 | 1 |
| #042 | 1.46 | 0 | 1.54 | 0 |
| #030 | 1.48 | 0 | 1.55 | 0 |
| #034 | 3.26 | 0 | 3.34 | 0 |

Abbreviations: DOR, duration of response; EFS, event-free survival

**Supplemental Table 3:** EFS and DOR of the patients analyzed by spectral flow cytometry

| Cell Type | Subtype | Positive Markers (expressed) | Negative Markers  (not expressed) |
| --- | --- | --- | --- |
| T |  | CD3E, CD3G, CD3D, TRAC |  |
| CD4 | T | CD4 | CD8A, CD8B |
| CD4_Treg | CD4 | FOXP3 | EOMES, IFNG, GATA3 |
| CD4_Tr1 | CD4 | EOMES | FOXP3, IFNG, GATA3 |
| CD4_Th1/17 | CD4 | IFNG | FOXP3, EOMES, GATA3 |
| CD4_Th2 | CD4 | GATA3 | FOXP3, EOMES, IFNG |
| CD8 | T | CD8A, CD8B | CD4 |
| CD8_exhausted | CD8 | ENTPD1, LAG3, HAVCR2 |  |
| CD8_resident_memory | CD8 | ZNF683, ITGAE | ENTPD1, LAG3, HAVCR2 |
| CD8_early_memory | CD8 | CD28, EOMES, GZMK | ENTPD1, LAG3, HAVCR2, ITGAE |
| NK |  | KLRD1, GNLY, NKG7 | CD4, CD8A, CD8B, CD3E, CD3G, CD3D |
| NKT | T | KLRD1, GNLY, NKG7 | CD4, CD8A, CD8B |

**Supplemental Table 4:** Markers used in the Garnett-classifier training phase

| Fluorophore | Marker | Clone | Source |
| --- | --- | --- | --- |
| BUV395 | CCR7 | 2-L1-A | BD |
| BUV496 | CD45RA | 5H9 | BD |
| BV421 | CD84 | 2G7 | BD |
| V500 | CD45 | 2D1 | BD |
| BV605 | CD62L | DREG-256 (562719) | Biolegend |
| BV650 | CD27 | L128 | BD |
| BV711 | TIGIT | A15153G | Biolegend |
| BV750 | CD127 | HIL-7R-M21 | BD |
| BV786 | CD3 | SK7 (563800) | BD |
| VB515 | CD19 CAR | REA746 | Miltenyi |
| BB700 | BTLA | 746166 | BD |
| PE | PD-1 | EH12.1 | BD |
| PE-CF594 | CD8 | RPA-T8 | BD |
| PECy7 | CD117 | 104D2 | BD |
| APC | TGFBR2 | W17055E | Biolegend |
| Alexa 647 | TIM3 | 7D3 | BD |
| APC-R700 | CD56 | NCAM16.2 (657886) | BD |
| APC-Cy7 | CD25 | M-A251 | Biolegend |
| APC/Fire 810 | CD4 | SK3 | BD |
| BUV661 | CD15 | W6D3 | BD |
| PB | HLA-DR | L243 | Biolegend |
| PEcy5 | CD33 | WM53 | Biolegend |
| BV570 | CD14 | M5E2 | Biolegend |
| BUV737 | TLR2 (CD282) | 11G7 | BD |
| BUV805 | CD11b | D12 | BD |
| BV480 | CD226 | 11A8 | BD |
| BUV615 | LAG3 (CD223) | T47-530 | BD |
| BUV 563 | CD16 | 3G8 | BD |
| Zombie NIR | Live cells |  | Biolegend |

**Supplemental Table 5:** Flow cytometry 30 color panel for validation

| Fluorophore | Marker | Clone | Source |
| --- | --- | --- | --- |
| FITC | CD6 | BL-CD6 | Biolegend |
| BV786 | CD3 | SK7 (563800) | BD |
| V500 | CD45 | 2D1 | BD |
| PE-CF594 | CD8 | RPA-T8 | BD |
| APC-H7 | CD4 | SK3 | BD |
| BV605 | CCR2(CD192) | K036C2 | Biolegend |

**Supplemental Table 6:** Flow cytometry 7 color panel for validation

| Fluorophore | Marker | Clone | Source |
| --- | --- | --- | --- |
| Zombie NIR | Live cells |  | Biolegend |
| Zombie Aqua | Live cells |  | Biolegend |
| APC/Fire 810 | CD4 | SK3 | BD |
| BUV496 | CD45RA | 5H9 | BD |
| APC-Cy7 | CD25 | M-A251 | Biolegend |
| V500 | CD45 | 2D1 | BD |
| BV605 | CD62L | DREG-256 (562719) | Biolegend |
| BV650 | CD27 | L128 | BD |
| BV711 | TIGIT | A15153G | Biolegend |
| BV750 | CD127 | HIL-7R-M21 | BD |
| BV786 | CD3 | SK7 (563800) | BD |
| VB515 | CD19 CAR | REA746 | Miltenyi |
| APC | CD10  TGFBR2 | 746166 | BD |
| PE | PD-1 | EH12.1 | BD |
| PE-CF594 | CD8 | RPA-T8 | BD |
| APCCy7 | CD19 | HIB19 | BD |
| PECy7 | CD19 | HIB19 | BD |
| BV480 | CD226 | 11A8 | BD |
| BV615 | LAG3 | T47-530 | BD |
| Alexa 647 | TIM3 | 7D3 | BD |

**Supplemental Table 7:** Flow cytometry 18 color panel for hypoxic co-colture validation

| Fluorophore | Marker | Clone | Source |
| --- | --- | --- | --- |
| Zombie NIR | Live cells |  | Biolegend |
| V450 (PB) | hCD45 | HI30 | Invitrogen |
| PerCP | mCD45 | 30-F11 | Biolegend |
| BV711 | CD33 | WM53 | Biolegend |
| PE | CD19 | SJ21C1 | eBiosciences |
| APC | CD3 | OKT3 | Biolegend |
| APC | TGFBRII | W17055E | Biolegend |
| FITC | CD19 | SJ21C1 | Biolegend |
| AF700 | CD8 | SK1 | Biolegend |
| BV605 | CD62L | MEL-14 | Biolegend |
| PE | CD45RA | HI100 | Biolegend |
| BV786 | CD3 | UCHT1 | BD Bioscience |
| BV711 | TIGIT | VSTM3 | Biolegend |
| APC-Cy7 | PD1 | HB3 | Invitrogen |
| PECy7 | aBiot CAR | REA746 | Miltenyi |
| Biotin | CD19 |  | Miltenyi |
| BV 510 | TLR2 | 11G7 | BD Bioscience |
| V450 (PB) | HLA-DR | I243 | Biolegend |
| APC | hCD45 | HI30 | Biolegend |
| FITC | CD19 | SJ25C1 | Biolegend |
| AF700 | hCD17 | 104D2 | Biolegend |
| BV605 | CCR2 | K036C2 | Biolegend |
| PE | CD33 | WM53 | Biolegend |
| BV786 | CD11b | ICRF44 | BD Bioscience |
| BV711 | CD14 | MOP9 | BD Bioscience |
| BV711 | CD15 | W6D3 | BD Bioscience |
| APC-Cy7 | CD34 | 581 | Biolegend |
| PECy7 | VEGFR2 | A16085H | Biolegend |

**Supplemental Table 8:** Flow cytometry antibodies list for in vivo staining

**Supplemental Figures:**

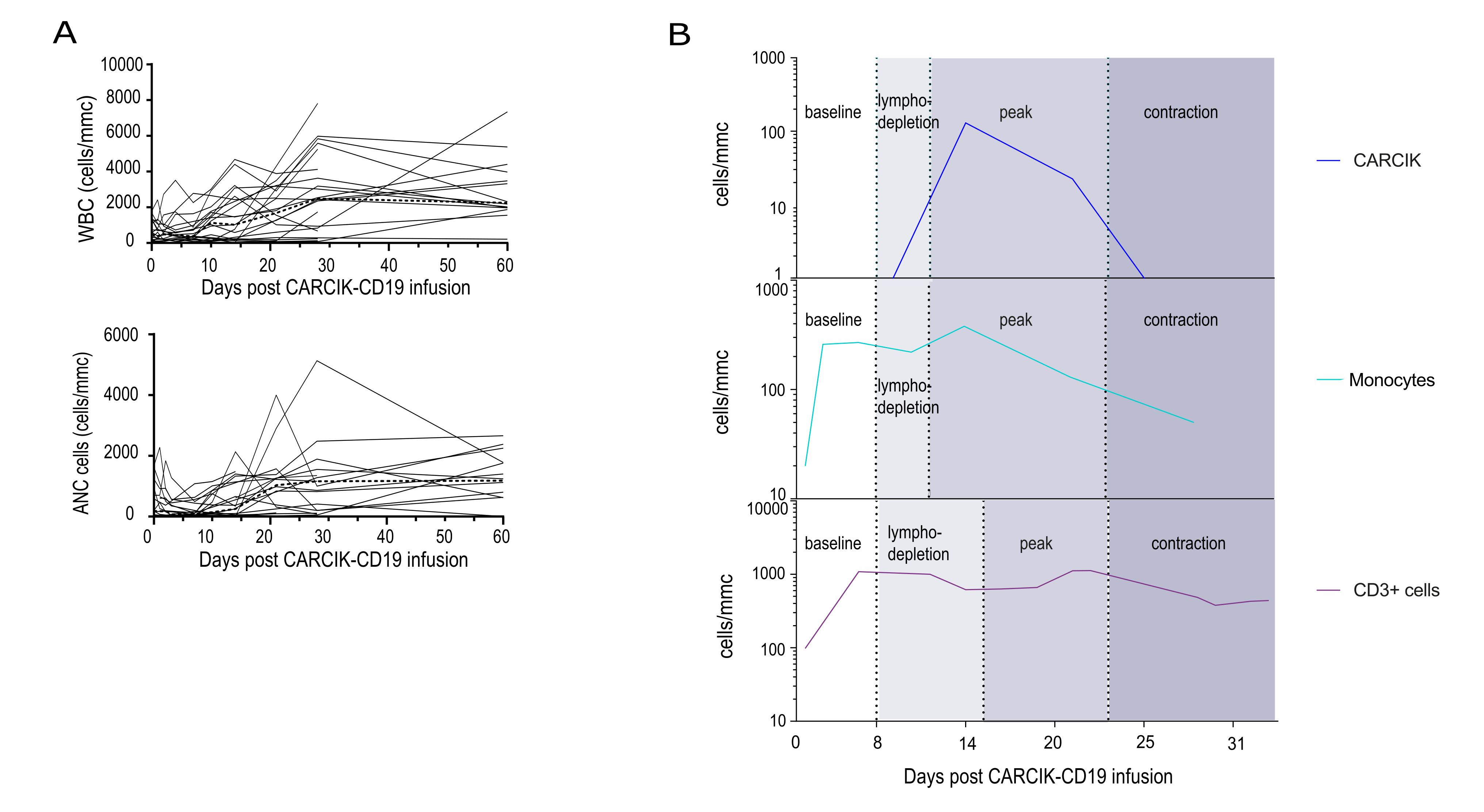

**Supplemental Figure 1.** **WBC and ANC kinetics after CARCIK-cell infusion.** (A) WBC, and ANC in 21 patients treated with allogeneic CARCIK-CD19. The median is depicted as a dashed line. (B) CAR T cells, monocytes and CD3+ cell cellular dynamics of a representative adult patient with B-ALL treated with CARCIK-CD19 showing a biphasic kinetic due to lymphodepletion, expansion and contraction.

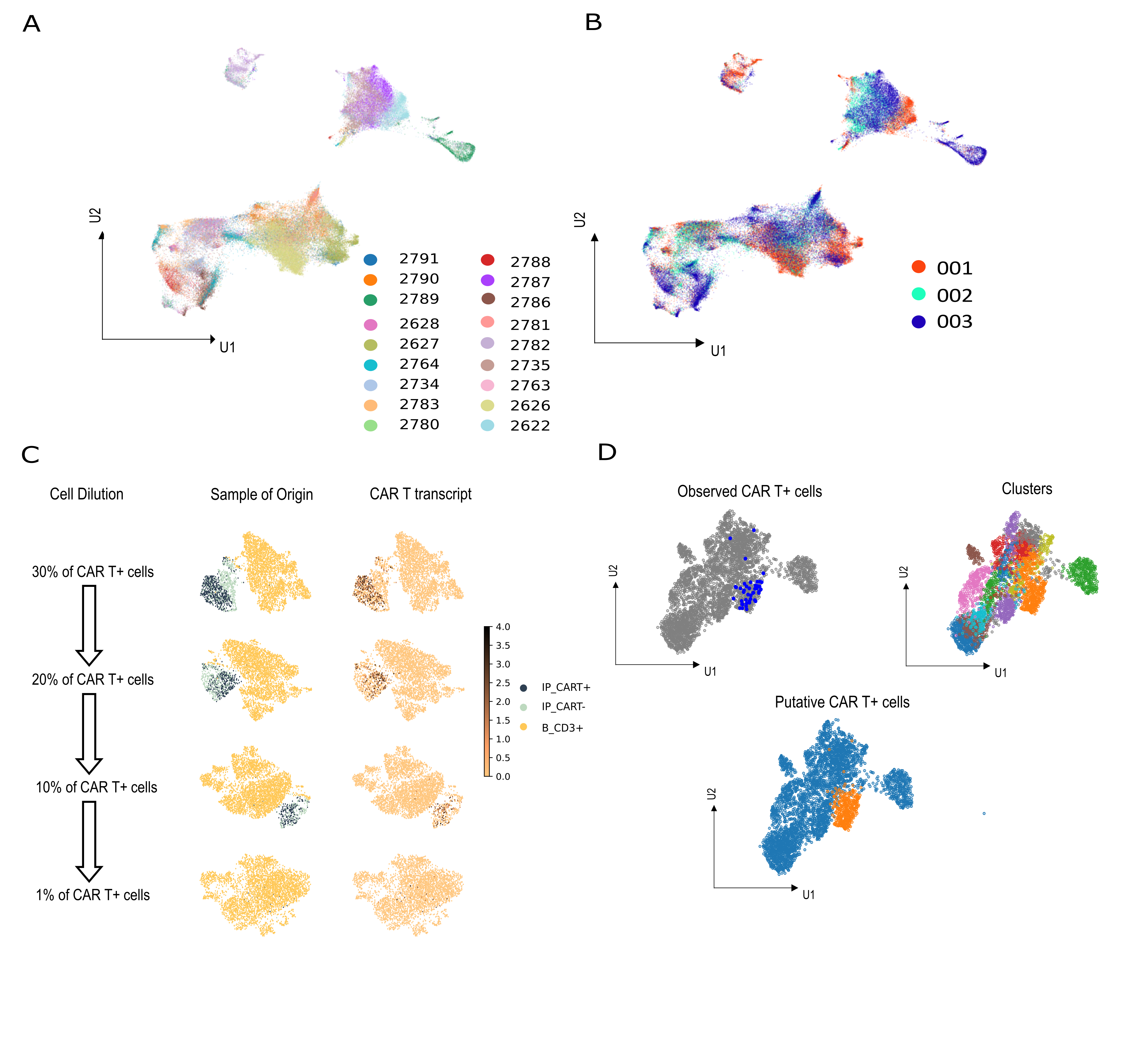
**Supplemental Figure 2.** **Single-cell RNA sequencing of the BM following CD19 CAR T cell therapy.** (A-B) UMAP embedding of multidimensional scRNA-seq data by libraries (left) and patient samples (right). (C) In silico limiting dilution assay for evaluating the sensitivity of detection of the CAR transcript in complex cellular mixtures. The reads form the IP were diluted with reads from BM pre-infusion (CD45+ CD3+ fraction). Limiting concentration of CAR T cells to yield good separation was between 5% and 1%. (D) CAR gene expression is depicted in the UMAP space (left). High-resolution clustering of CD3+ cells (right) allows the recognition of a qualitatively enriched cluster in CAR T cells that was annotated as putative CAR+ (bottom).

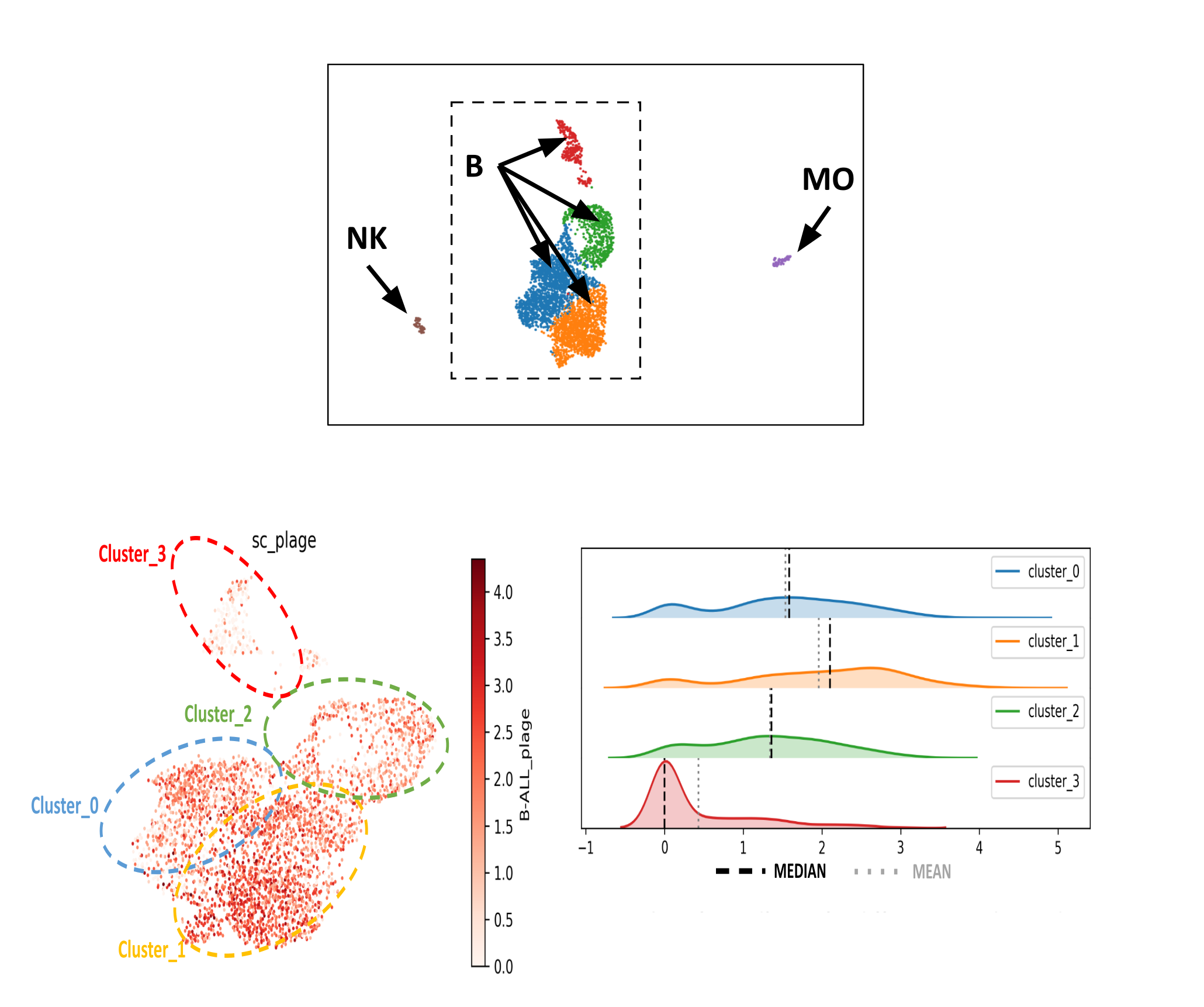

**Supplemental Figure 3.** **Identification of the malignant and healthy B-cell clusters.** High-resolution clustering of the CD45+CD3- libraries and application to the four B-cell clusters of a gene signature discriminating leukemic blasts from healthy cells.

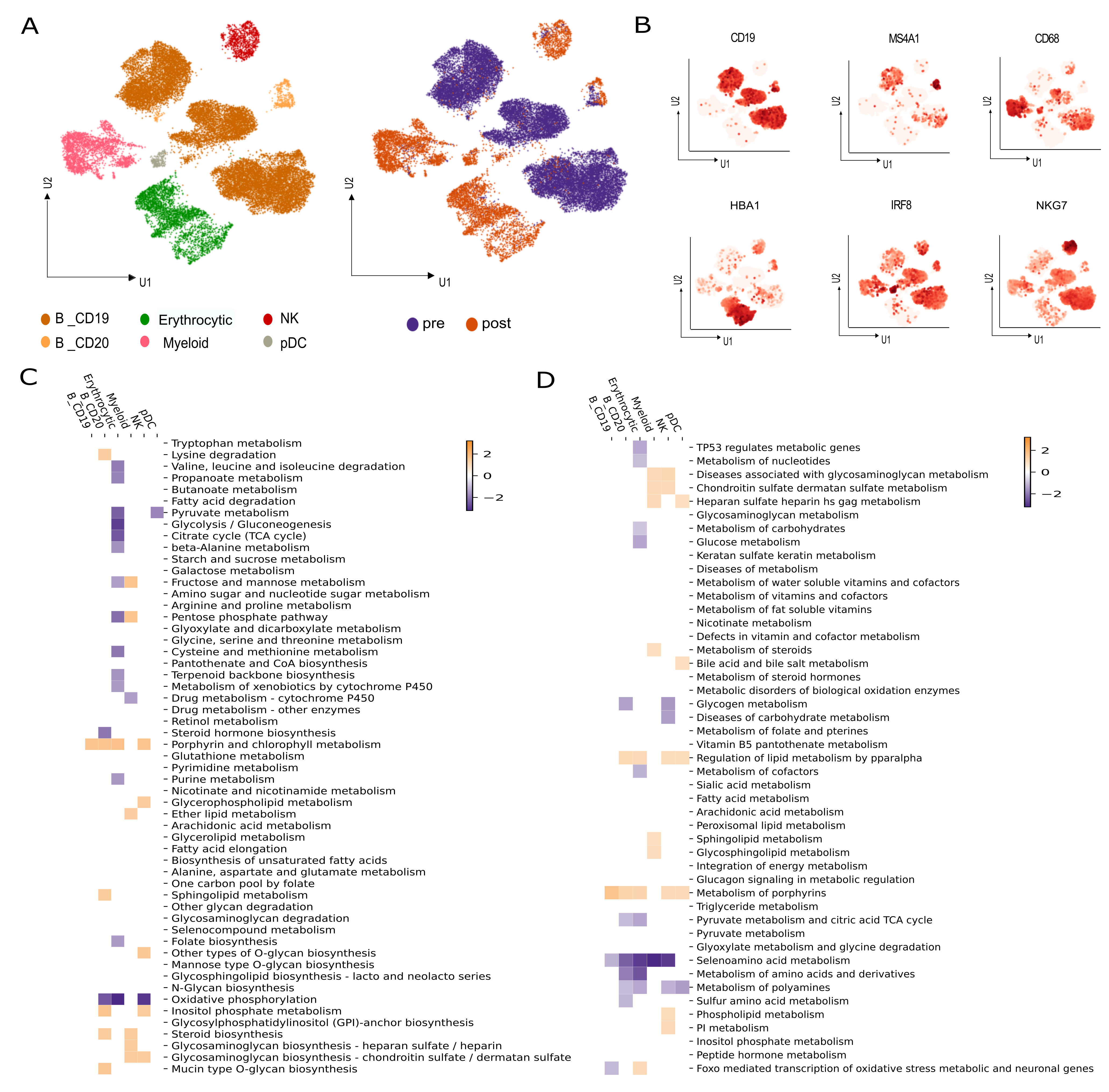

**Supplemental Figure 4.** **CD45+CD3- compartment analysis in CAR T-cell treated samples versus untreated.** (A) UMAP visualization of the CD45+CD3- libraries as cell clusters (left) and time points (right). (B) Marker-based cell type identification analysis allowed prediction of six broad immune cell types across all profiled single cells in CD45+CD3- fraction. (C-D) Significantly enriched KEGG (left) and Reactome (right) gene sets in post-treatment versus pre-treatment samples within each of the immune cell types.

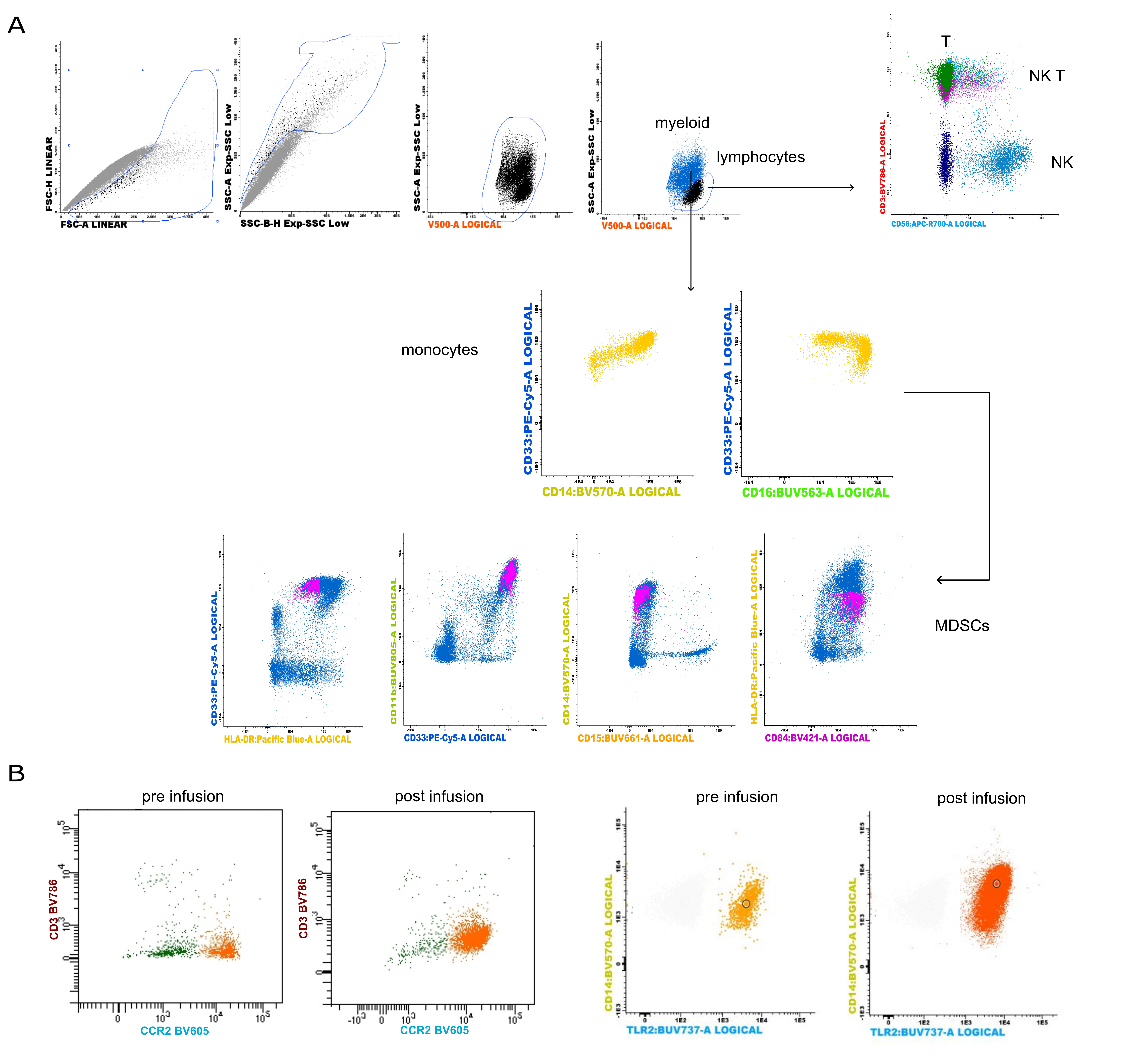

**Supplemental Figure 5. Flow cytometry analysis with Infinicyt for the identification of the myeloid cell subset.** (A) Representative gating strategy to remove debris, and identify, within the lymphocytes’ gate, T cells (CD3+), NKT cells (CD3+CD56+), NK cells (CD3-CD56+), and, within the myeloid gate, monocytes (CD33+ CD14+HLA-DR+CD16+), and MDSCs (CD11b+CD33+HLA-DR-/low CD14-CD15+ or CD14+CD15-). (B) Identification of populations expressing CCR2 and TLR2 markers in the myeloid compartment.

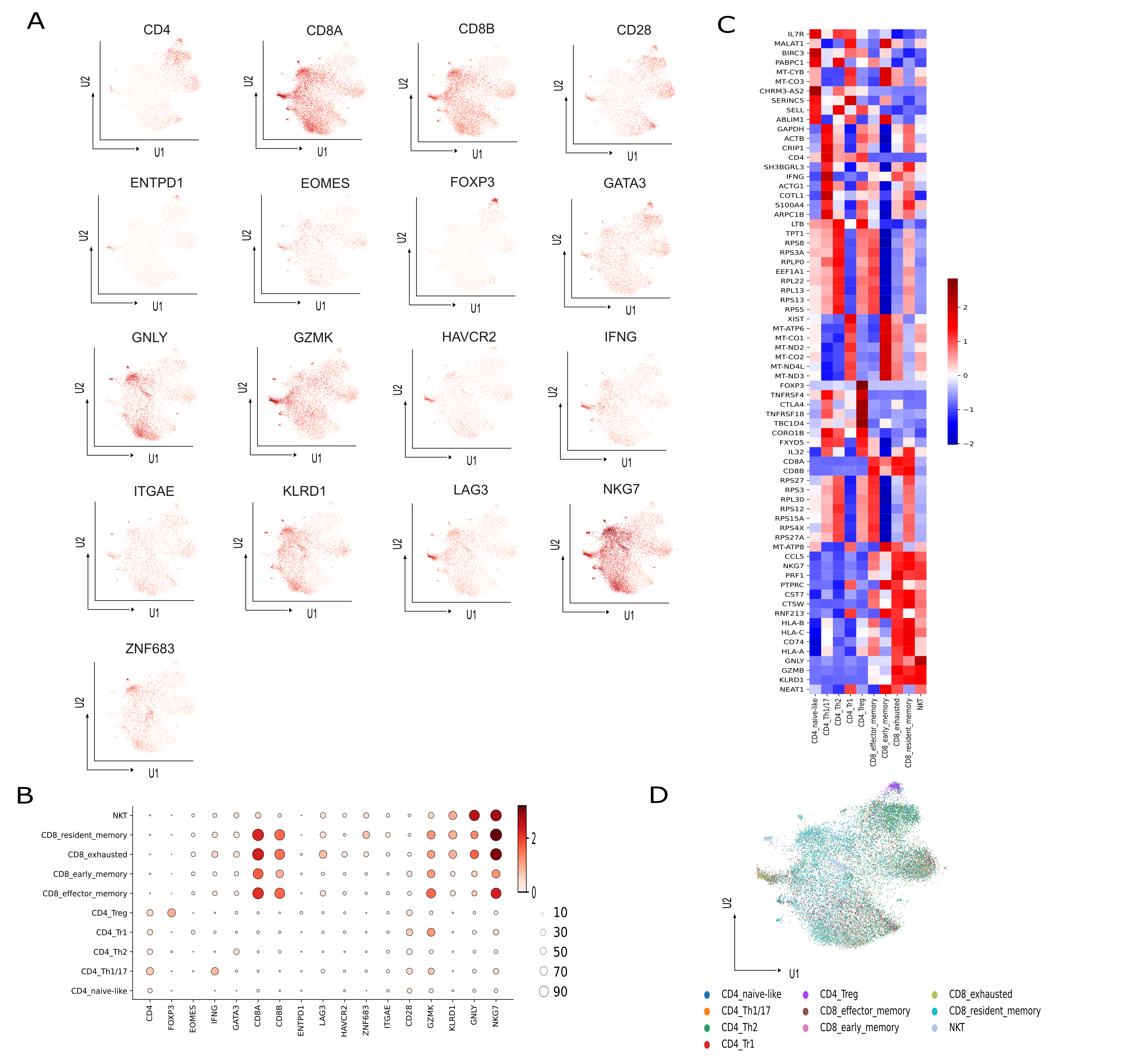

**Supplemental Figure 6. Identification of bone marrow CD3+ compartments.** (A) Expression levels of top CD3+ cluster-specific markers overlaid on the UMAP. (B) Dot Plot of quantitative related markers expression in CD45+CD3+ cell clusters. (C) Gene expression heatmap of the top ten DEG between CD45+CD3+ cell clusters. (D) UMAP projection of identified cell clusters, including naive-like CD4+, CD4+ Th1/17, CD4+ Th2, CD4+ Tr1, CD4+ Treg, effector memory CD8+, early memory CD8+ , exhausted CD8+, resident memory CD8+ and NKT cells.

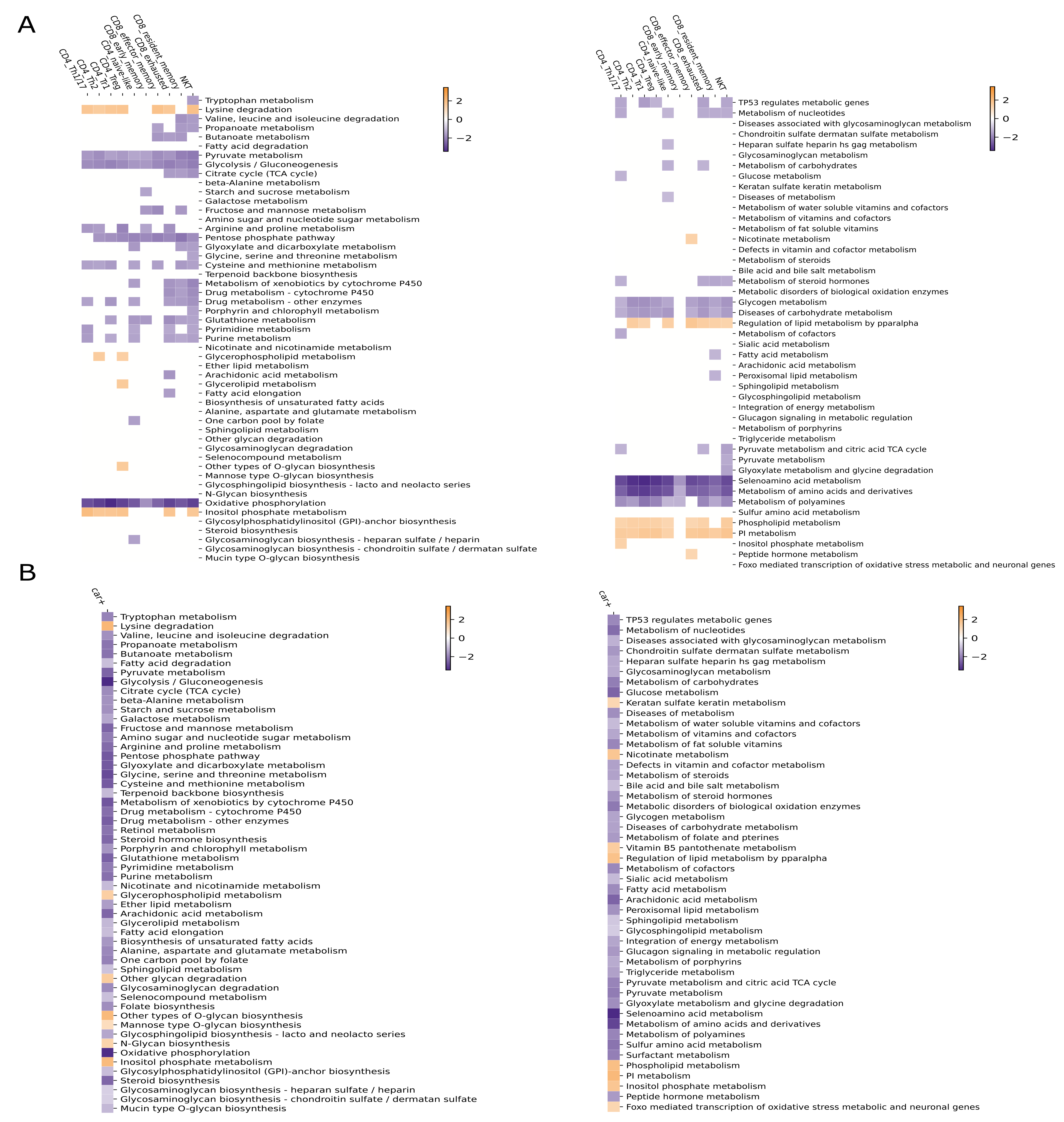

**Supplemental Figure 7. Endogenous T cells and infused CAR T cells showed increased lipid metabolism post treatment.** (A-B) Significantly enriched KEGG (left) and Reactome (righ) gene sets in post–treatment versus pre-treatment samples within each of the A) endogenous CD3+ immune cell types (top) and B) within CAR T cells (below).

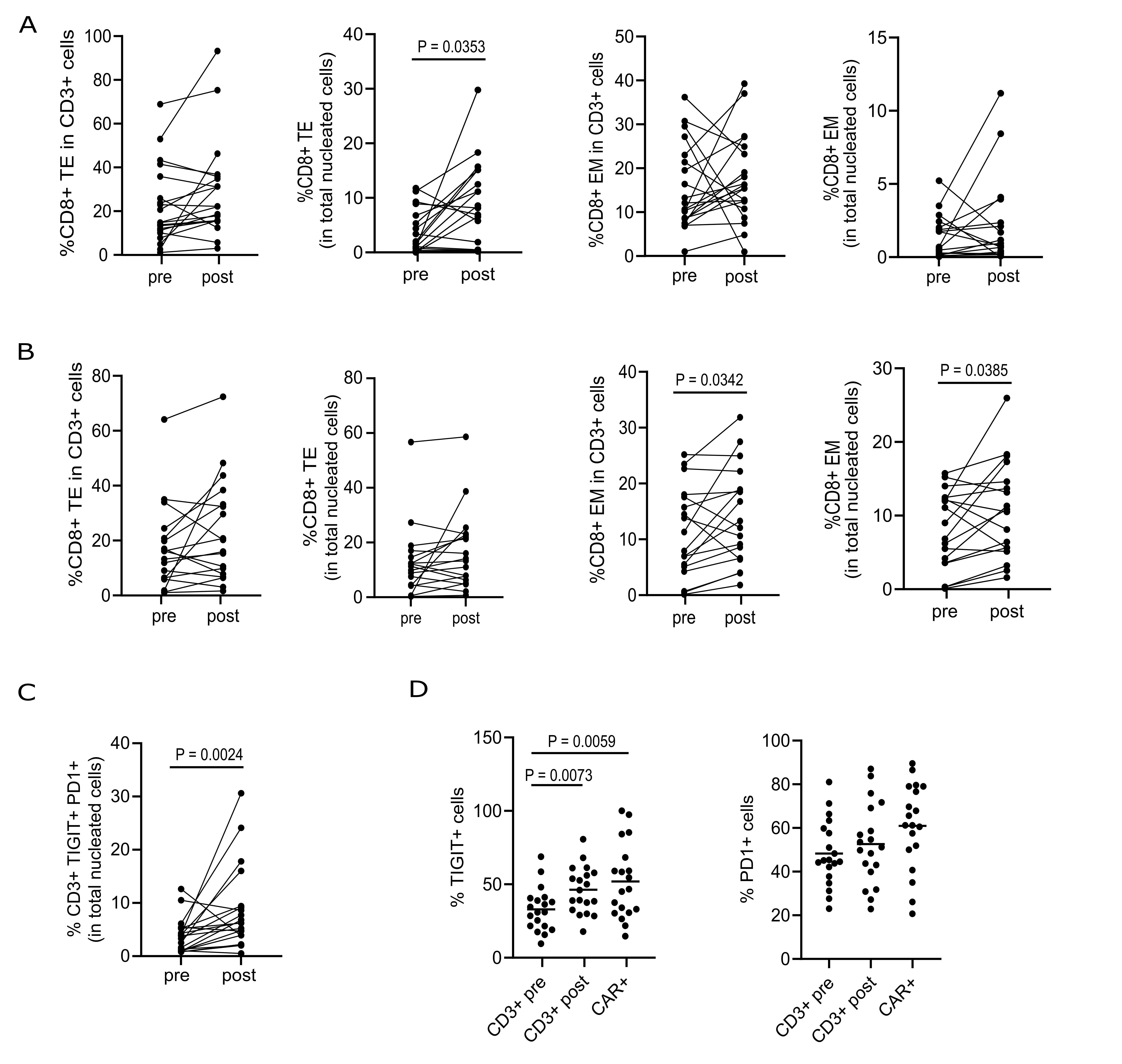

**Supplemental Figure 8. Comprehensive analysis of the T-cell compartments by spectral flow cytometry.** (A) Flow cytometry analysis (n=20 pre- and n=20 post- treatment paired BM samples) with Infinicyt depicting the frequencies of terminal effector CD8+ cells in CD3+ cells and total nucleated cells and of effector CD8+ in CD3+ cells and total nucleated cells. Wilcoxon matched-pair t test. *, P < 0.05. (B) FlowSOM-generated analysis depicting the frequencies of terminal effector CD8+ cells in CD3+ cells and total nucleated cells and of effector CD8+ in CD3+ cells and total nucleated cells. Wilcoxon matched-pair t test. *, P < 0.05. (C) Flow cytometry analysis with Infinicyt depicting the frequencies of TIGIT+PD1+CD3+ cells in total nucleated cells. Wilcoxon matched-pairs t test. **, P < 0.01. (D) Flow cytometry analysis with Infinicyt depicting the frequencies of TIGIT+ CD3+ and PD1+ CD3+ cells in total CD3+ cells before and after CAR T-cell treatment and in BM-retrieved CAR T cells. One-way ANOVA with Turkey multiple comparison **, P < 0.01

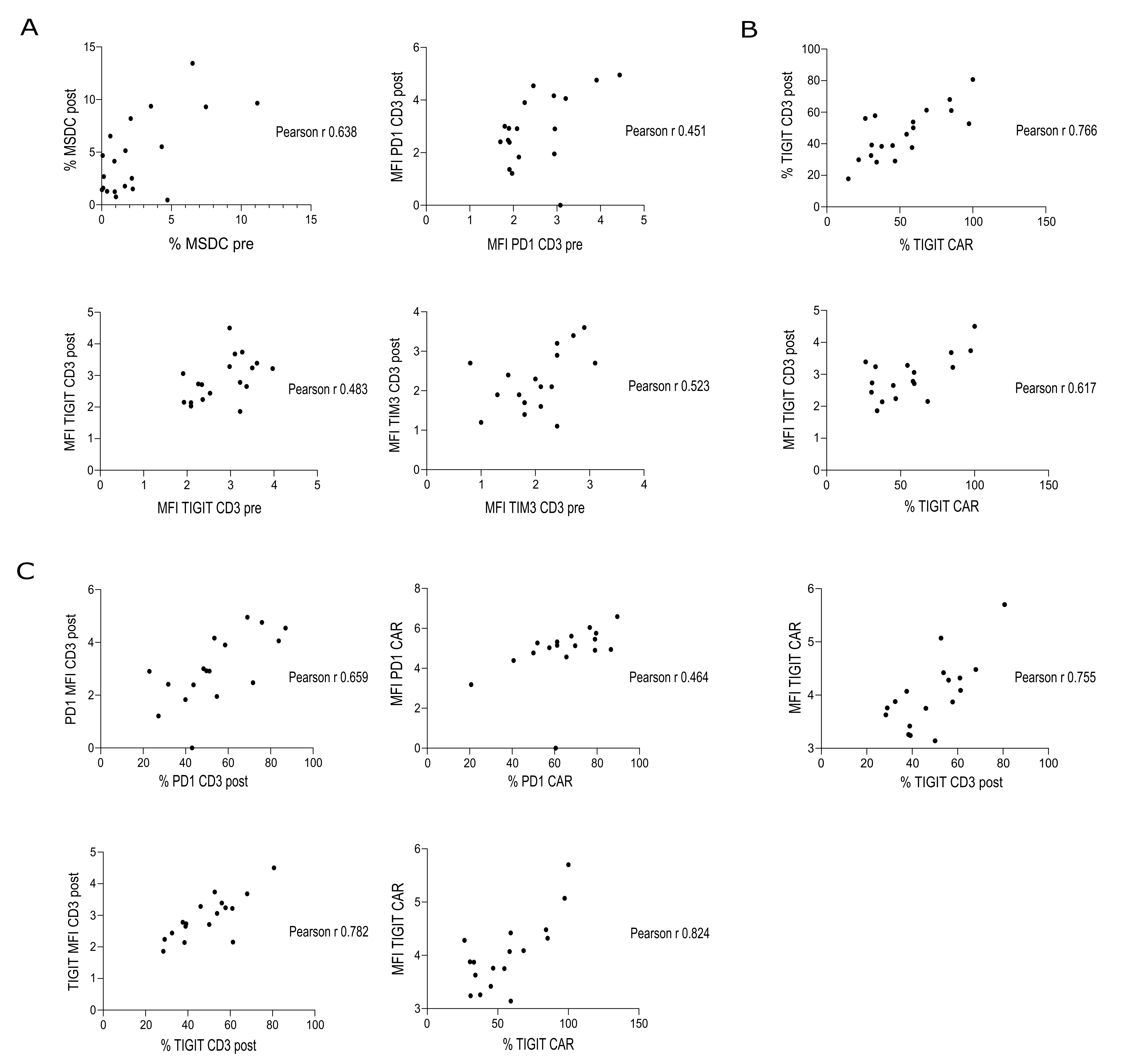

**Supplemental Figure 9. Pearson correlation between features associated with TME remodeling. (A)** Correlation between the frequency of MDSCs and PD1, TIGIT, and TIM3 MFI in endogenous T cells before and after treatment. **(B)** Correlation between the frequency / MFI of TIGIT in endogenous T cells and those in CAR T cells after treatment. **(C)** Correlation between the frequency and the MFI of PD1 and TIGIT in endogenous T cells and CAR T cells before and after treatment.

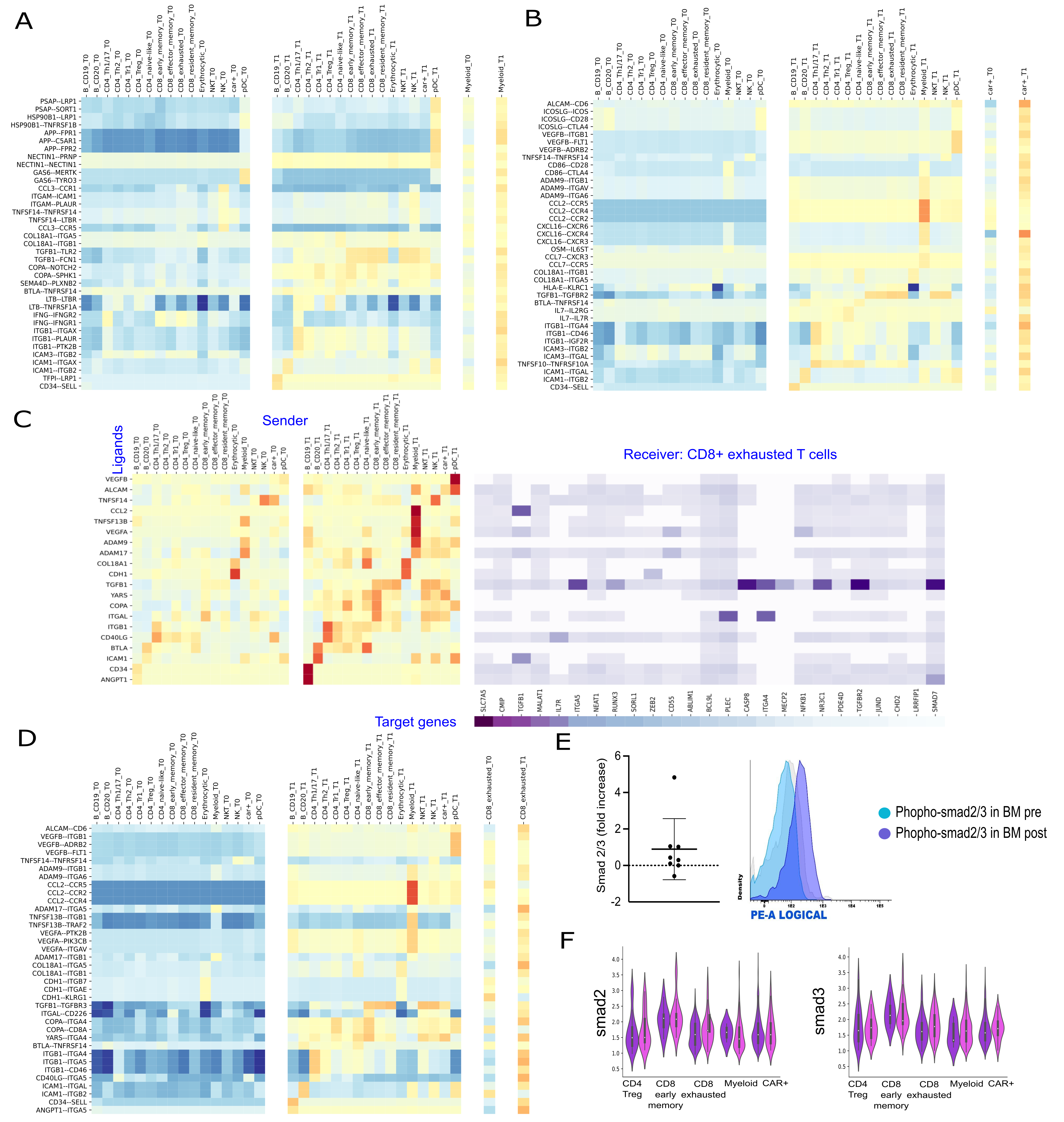
**Supplemental Figure 10. NicheNet-inferred ligand-receptor interactions revealed an autocrine TGF-β signaling loop in CAR T cells and endogenous T cells.** (A-B) NicheNet Heatmap depicting potential ligand-receptor link activities assessed using A) all niche as sender and myeloid compartment as receiver and B) using all niche as sender and CAR T cells as receivers. (C) NicheNet’s ligand activity prediction (left) and ligand-target matrix (right) assessed using all niche as sender and CD8+ exhausted T cells as receiver. (D) NicheNet Heatmap depicting potential ligand-receptor link activities assessed using all niche as sender and CD8+ exhausted T cells as receiver. (E) Phospho-flow analysis of smad2/3 phosphorylation in CD3+ cells assessed by intracellular staining. Data illustrate the mean ± SD of the fold increase expression in BM pre-treatment compared to post-treatment. One representative patient of results in 8 paired samples is shown. (F) Violin plots of scRNA-seq data depicting smad2 and smad3 expression distribution in different immune cell clusters before and after treatment

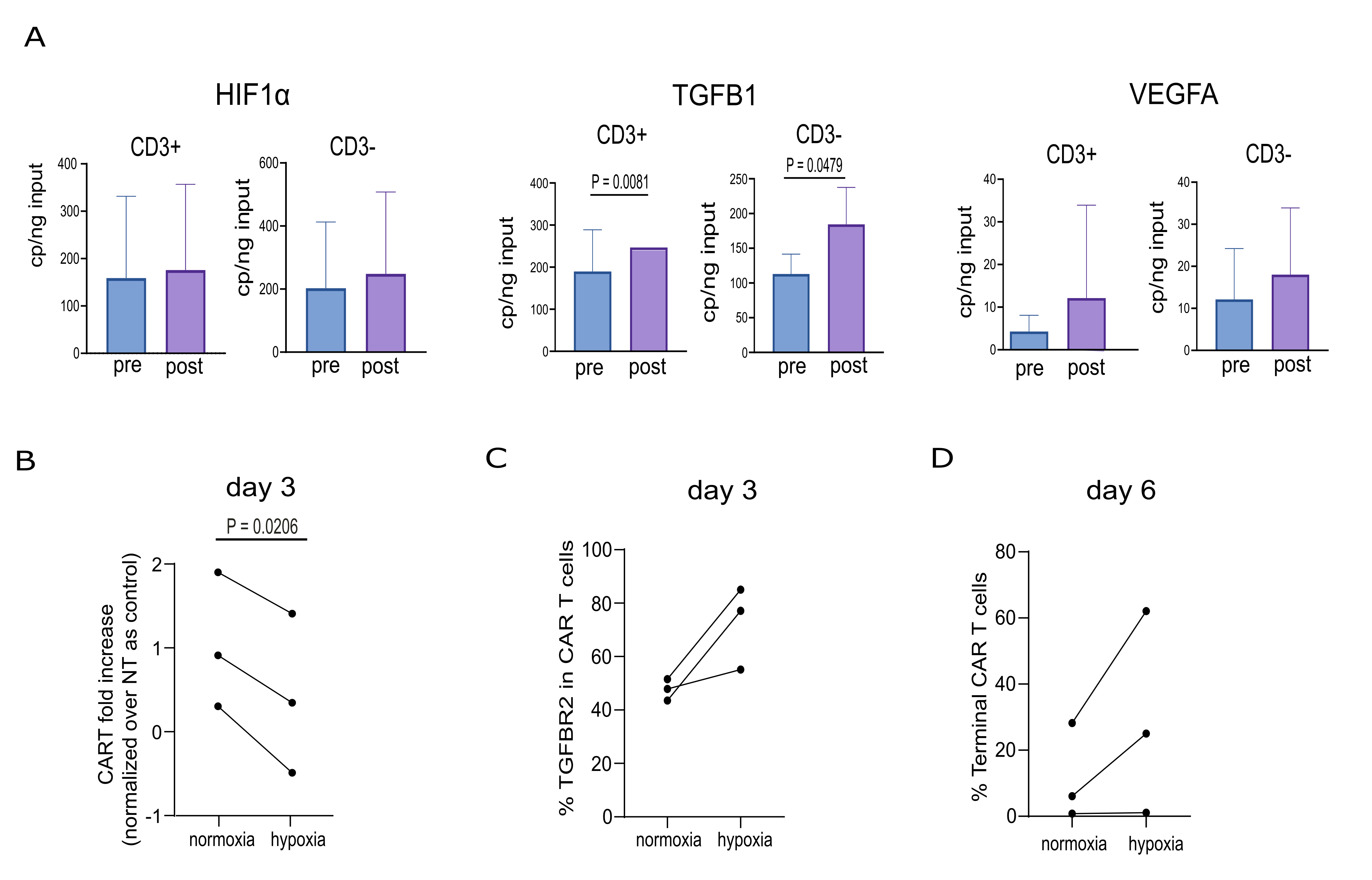
**Supplemental Figure 11. Chronic antigen stimulation in hypoxic conditions promotes T-cell dysfunction.** (A) ddPCR validation of HIF1α, TGFB1 and VEGFA genes in purified CD3+ and CD3- cell subsets (n=13 paired pre- and post- BM treatment samples). Data illustrate the mean ± SD. Wilcoxon matched-pair t test. (B) Flow cytometry analysis depicting CAR T-cell fold increase in hypoxic compared to normoxic condition after 3 days of co-culture. (C) Flow cytometry analysis depicting the frequencies of TGFBR2+ CD3+ cells after 3 days of co-culture. (D) Flow cytometry analysis depicting the frequencies of terminal CAR+ cells after 6 days of co-culture. Data illustrate the mean ± SD of triplicates from three different donors. Wilcoxon matched-pair t test. *, P < 0.05.

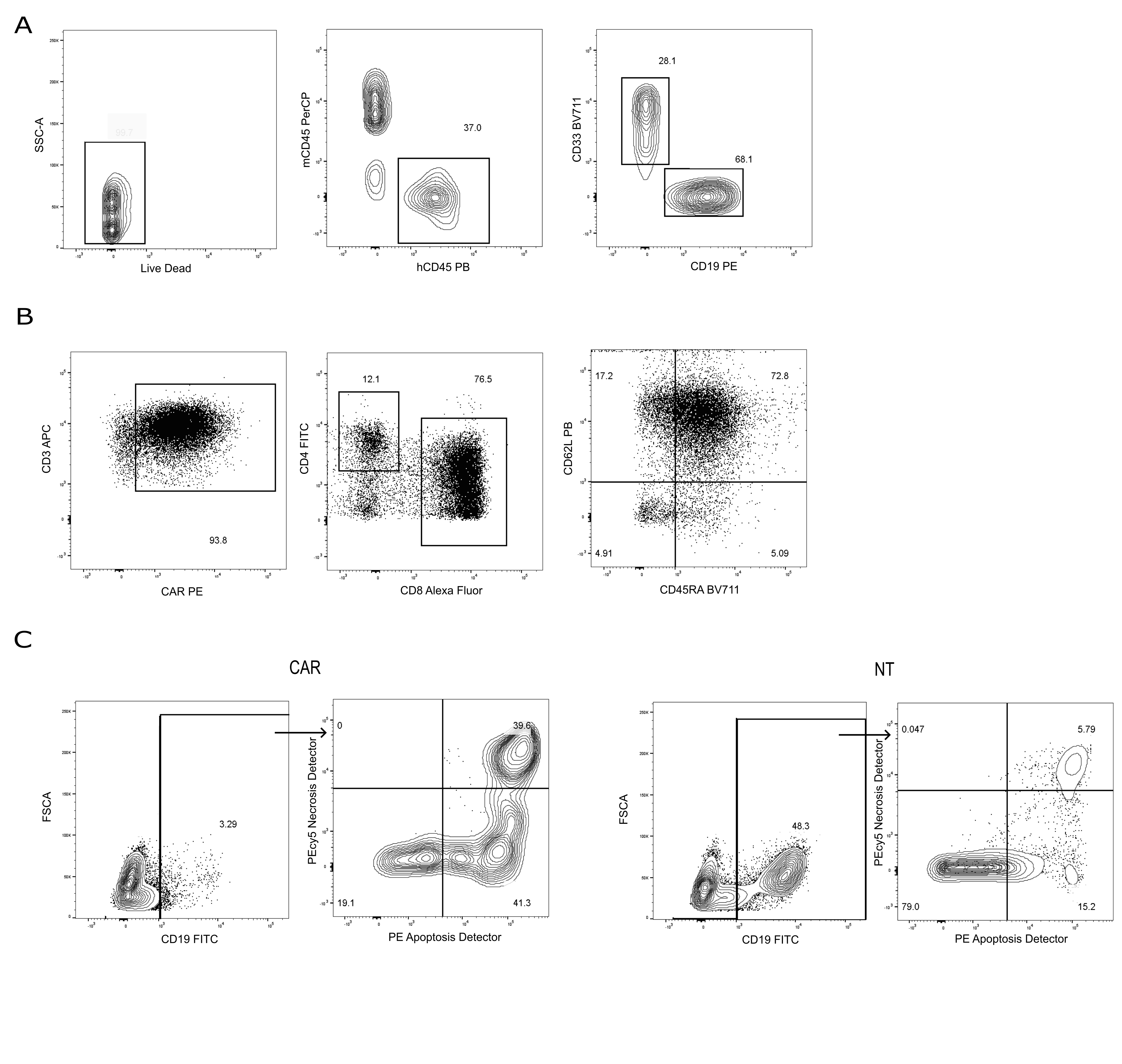
**Supplemental Figure 12. CAR T cells generated from HSPC-humanized mice demonstrate efficient killing in vitro.** (A) Representative Flow cytometry immunophenotyping of the PB of humanized animals engrafted with human CD34+ cells. (B) Phenotype of CAR T cells generated from splenocytes harvested from mice humanized with the same HSPC donor by lentiviral transduction of an anti-CD19CAR.BBz. (C) Killing activity of CAR T cells generated from HSPC-humanized mice after 24 hours of co-culture with Nalm6 at an E:T ratio of 1:1. Representative dot plots corresponding to one of the triplicate showing target cell apoptosis and necrosis.

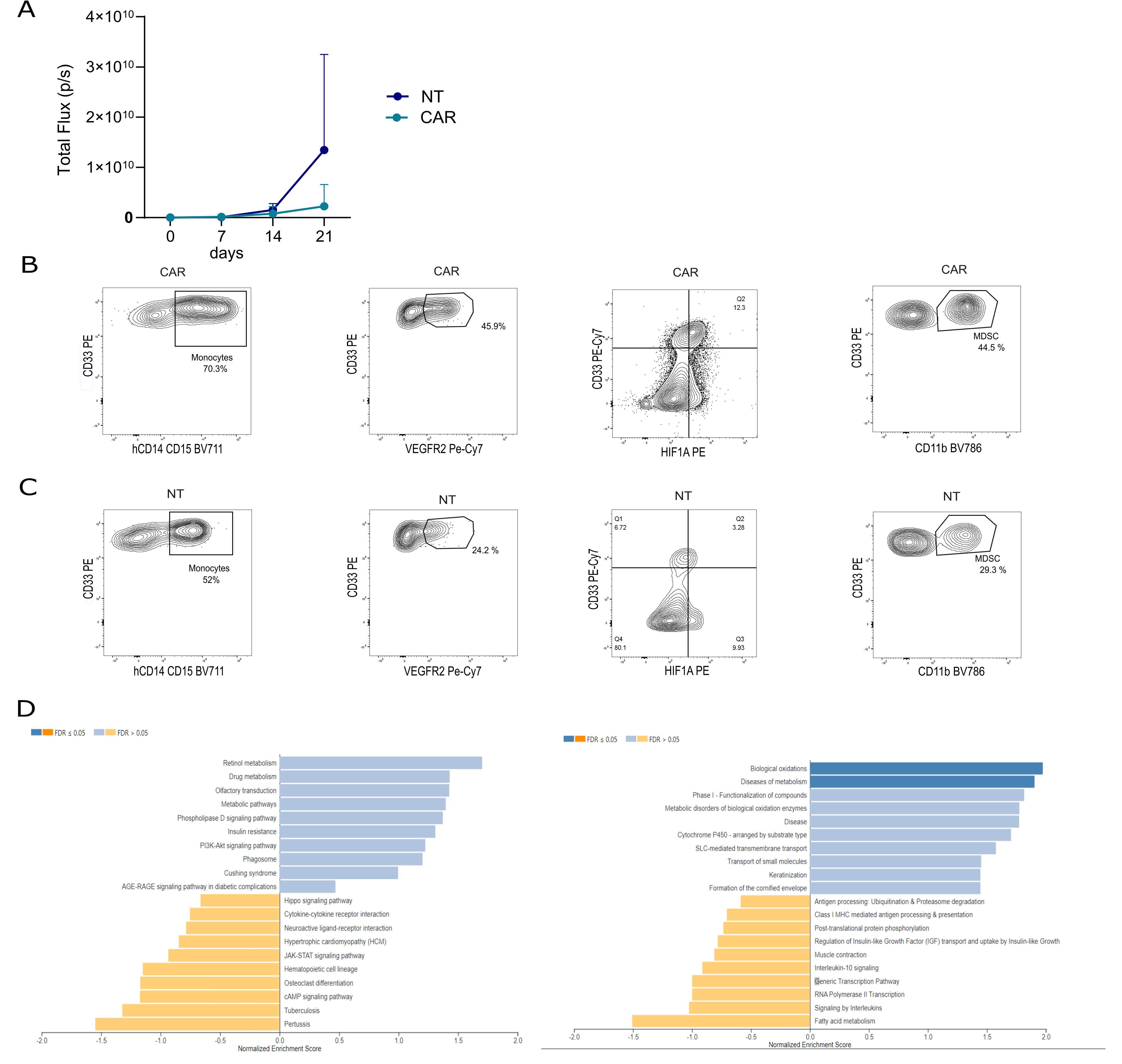
**Supplemental Figure 13. Remodeling of BM microenvironment after CAR T infusion in tumor-bearing humanized mice.** (A) Bioluminescence intensity (total flux per second) over time in mice receiving the indicated treatment. Two-way ANOVA with Turkey multiple comparison (*compared with NT). (B-C) Representative flow cytometry dot plot of monocytes, VEGFR2+CD33+ cells, HIF-1α^high^CD33+ cells and MDSC in BM of mice treated with CAR T or NT cells. One representative mouse is shown. (D) Significantly-enriched KEGG (left) and Reactome (right) gene sets in purified myeloid cells of mice treated with CAR T or NT cells, FDR (-logpvaladj) < 0.05.
